# Commensal bacteria can inhibit the growth of *P. aeruginosa* in cystic fibrosis airway infections through a released metabolite

**DOI:** 10.1101/2023.02.03.526996

**Authors:** Andrew Tony-Odigie, Alexander H. Dalpke, Sébastien Boutin, Buqing Yi

**Author notes:** Senior author (Sébastien Boutin, Buqing Yi). **Correspondence:** Dr. Buqing Yi, Institute of Medical Microbiology and Virology, University Hospital Carl Gustav Carus, Dresden, Saxony, Germany. Fetscherstraße 74, 01307 Dresden, Germany.

## Abstract

In cystic fibrosis (CF), infections with *Pseudomonas aeruginosa* or other typical pathogens play a critical role in eliciting disease progression, leading to tissue damage and finally loss of lung function. Previous observations showed that the presence of various commensal bacteria and a higher airway microbiome diversity were associated with better lung function and less severe disease burden. Thus, the hypothesis was raised that commensal bacteria might be able to interfere with pathogenic bacteria. In this study, we aimed to identify airway commensal bacteria that inhibit the growth of *P. aeruginosa*.

Through a screening experiment of co-culture with *P. aeruginosa* PAO1, we could identify more than 30 CF commensal strains from various species that inhibited the growth of *P. aeruginosa*. With multiple selected strains, we further verified the results with *P. aeruginosa* CF isolates and several other pathogens isolated from CF patients, and most of the identified commensal strains showed consistent results strongly inhibiting the growth of diverse CF pathogens.

The underlying mechanisms of the growth-inhibition effects were first investigated through genomic analysis by comparing strains with and without growth-inhibition effects, which revealed that genes responsible for carbohydrate transport and metabolism were highly enriched in the inhibitory commensals. Metabolite analysis and functional analysis showed that commensals with inhibitory effects produce large amounts of acetate. Exogenous addition of acetate under a low pH inhibited the growth of *P. aeruginosa*, indicating acetate produced and released by commensals may affect the growth of *P. aeruginosa* living in the same microenvironment.

In summary, through co-culture of *P. aeruginosa* with commensals, we could identify that a variety of airway commensal strains can inhibit the growth of *P. aeruginosa* by producing acetate. The data provide insights into possible novel strategies for controlling infections in people with CF and also emphasize the importance of preserving airway commensals when designing infection treatment strategies.

## Introduction

Cystic fibrosis (CF) is the most common fatal hereditary lung disease ^1^. CF pathology nowadays is typically characterized by a progressive loss of lung function through cycles of infection, hyper-inflammation and tissue damage, and infections with *Pseudomonas aeruginosa* or other typical CF pathogens play a critical role in initiating this cycle ^1–3^. Although many people with CF (pwCF) are now treated with CFTR modulator therapy, results from modulator therapy studies indicate that this therapy itself can hardly eradicate chronic *P. aeruginosa* infection ^4–6^, and it is noteworthy that there are still around 20-30% of pwCF who cannot be treated with modulator therapy^7^. Thus, controlling chronic CF airway infections is still one important task for physicians in the healthcare management of pwCF ^4,8^.

To control respiratory bacterial infections, antibiotic treatment is widely used through inhalation, or oral/intravenous administration ^9,10^. Increased numbers of antibiotics prescribed to the patients are associated with significant decreases in microbial diversity, thus affecting local micro-ecology ^9,11^. Multiple CF studies originating from airway microbiome sequencing indicate that the airway microbiome diversity, in particular, the presence of various commensal bacteria e.g. Streptococci, is positively associated with better lung function and less severe disease burden ^11–14^. One hypothesis that has been put forward is that these commensal bacteria might be able to interfere with pathogenic bacteria by reducing or even preventing the harmful impacts of CF pathogens. In contrast, diminished microbial diversities in the airway (e.g. airways dominated by a certain CF pathogen) are often correlated with reductions in lung function ^15–21^. However, it is still unknown how to prevent the decrease of bacterial diversity in pwCF at an early stage. The complex interactions between pathogenic and commensal bacteria in the CF microbiome community are still underexplored, and the role of airway commensal microbes in the polymicrobial lung infection in pwCF remains unclear ^22–24^.

To investigate airway commensal-pathogen interactions, we performed a screening study with commensal bacterial strains isolated from airway samples of pwCF to identify strains that interfere with the growth of pathogenic bacteria and we aimed to identify underlying mechanisms.

## Methods and materials

### 1. Isolation of aerobic or facultative anaerobic microbes from CF sputum samples

As described previously ^25^, sputum samples of pwCF were streaked on Columbia Blood Agar (CBA) (Thermo Fisher Scientific, Waltham, USA) and cultured aerobically for 24h to isolate aerobic or facultative anaerobic strains. Isolated strains were identified by MALDI Biotyper System (Bruker Daltonik GmbH & Co. KG, Bremen, Germany), and then stored in skimmed milk at −20°C. Strains belonging to typical CF pathogen species were kept in the pathogen stock, while strains that are not typically considered classical pathogens and recognized as part of the commensal oropharyngeal microbiota based on literature information (e.g. ^26–30^) were kept in the commensal stock with details listed in sup. Table S1.

### 2. Screening study to identify commensals that may affect the growth of *P. aeruginosa* and other CF pathogens

A quantitative approach was employed using a genetically engineered *P. aeruginosa* strain, PAO1 pBPF-mcherry, producing red (via mCherry expression) fluorescent protein ^31^ to specifically monitor the growth of *P. aeruginosa* in co-culture (labelled as PAO1-mcherry). McFarland concentrations were measured using Columbia broth (CB) as growth medium. Each co-culture sample contained 4 μl McF0.5 concentration of PAO1-mcherry (approx. 7 x 10^5 CFUs) and 30 μl McF3.0 commensals (approx. 20-150 x 10^5 CFUs) in a final volume of 200 μl in 96 well plates. The 96-well plates were incubated and continuously recorded using a Synergy H1 Hybrid Multi-Mode Reader (BioTek Instruments, Inc., Vermont, USA) for 30h at 37°C and 5% CO2. Absorbance was read at 600nm to monitor the growth of bacteria, and the growth of *P. aeruginosa* was specifically measured by red fluorescence (RFU) which was recorded at 580nm (excitation) and 616nm (emission). For selected strains, CFUs were counted after plate streaking to further verify the results of RFU and OD600 readouts. Using the same experimental setting, we also tested multiple *P. aeruginosa* clinical isolates as well as other CF pathogenic isolates. *P. aeruginosa* CF clinical isolates were obtained from patients at different disease progression stages. All pathogenic strains used in this investigation are listed in sup. Table S2.

### 3. Whole genome sequencing of bacteria

Bacterial DNA was isolated with the DNeasy Blood & Tissue Kit (Qiagen, Hilden, Germany) following the manufacturer’s instructions. Library preparation and sequencing on a MiSeq Illumina platform (short-read sequencing 2×300bp) were performed as previously described ^32^. In short, raw sequences were controlled for quality using sickle (v1.33, parameter: -q 30 -l 45) and assembled with SPAdes 3.13.0 (with the option –careful and –only-assembler). Draft genomes were curated by removing contigs with a length <1000bp and/or coverage <10X. The quality of the final draft was controlled using Quast (v5.0.2). The draft genome sequences are deposited in the NCBI GenBank database under the project number PRJNA917188. Sequencing statistics and isolate data are available in the Supplementary material (sup. Table S3).

### 4. Whole-genome sequence analyses

Whole-genome sequence comparison by average nucleotide identity (ANI) was calculated for BLAST-based ANI (ANIb) with PYANI (Pritchard et al., 2016). For phylogenomic analysis of *Streptococcus* genomes, we used the sequence information of single-copy core genes present in 100% *Streptococcus* stains. The program MUSCLE ^33^ with default settings was used for creating the alignment of protein sequences. The alignments were cleaned up by removing positions that were gap characters in more than 50% of the sequences using trimAI ^34^. Sequence alignments of core genes were concatenated to give a single core alignment and a maximum likelihood phylogeny was then generated using the program ^35,36^ with a time-reversible aminoacid substitution model WAG, four gamma categories for rate heterogeneity and 1000 bootstrap replicates. The WAG model was chosen because it is applicable to phylogenetic studies of a broad range of protein sequences and generates more accurate phylogenetic tree estimates compared to many other models ^37^. Phylogenies were visualized using the Interactive Tree of Life ^38^.

### 5. Comparative functional genomic analysis

First, pangenome analysis was performed with anvi’o (version 7.1) ^39,40^ following the workflow for microbial pangenomics ^41^. In detail, PGAP ^42^ was used to identify open reading frames, then each database was populated with profile hidden Markov models (HMMs) by comparing to a collection of single-copy genes using HMMER ^43^. All genomes were annotated with the functional annotation source COGs (Clusters of Orthologous Genes) ^44,45^, which was accomplished by running anvi’o workflow “anvi-run-ncbi-cogs” that uses Diamond ^46^ to search the NCBI COG database and then annotates genomes accordingly. We used the Diamond software to calculate gene similarity and MCL ^47^ for clustering under the following settings: minbit, 0.5; mcl inflation, 2.0; and minimum occurrence, 1.

Second, as previously described ^48^, comparative functional enrichment analysis was performed by fitting a logistic regression (binomial Generalized Linear Model) to the occurrence of each gene function using group affiliation as the explanatory variable (“positive group” vs. “negative group”). Equality of proportions across group affiliation was tested using a Rao score test.

### 6. Profiling of short-chain fatty acids (SCFAs) in the conditioned medium of bacteria

Conditioned medium (CM) was prepared by introducing a Microbank bead (Pro-Lab Diagnostics, Ontario, Canada) of the microbial stock into a 12 ml propylene tube containing 5 ml of CB broth and incubating overnight at 37°C, 5% CO_2_ and 95% RH. 3 ml of this culture was then introduced into a 50 ml conical flask containing 17 ml of fresh CB and incubated for 48h. Afterwards, the culture was mixed, transferred into a 50 ml falcon tube, and centrifuged at 2,000 g for 10 mins. The supernatant was then filtered using a 0.22 μm filter. 20 μl of this CM was then inoculated onto a Columbia blood agar (CBA) plate and incubated overnight to confirm the absence of viable microbial cells. CM samples from a few selected commensal isolates as well as from PAO1-mcherry were sent to Lipidomix GmbH, Berlin, where their SCFAs profiles were analysed via high-performance liquid chromatography (HPLC).

### 7. Evaluation of the impact of SCFAs on the growth of *P. aeruginosa*

The evaluation was performed in a similar setting as for the co-culture’s test. Briefly, sodium acetate (Sigma-Aldrich, Taufkirchen, Germany) was added to CB to prepare defined concentrations. 4 μl McF0.5 concentration of PAO1-mcherry (approx. 7 x 10^5 CFU) or other *P. aeruginosa* strains were cultured in a final volume of 200 μl of CB (with or without SCFAs added) in 96 well plates. The pH values of CB were adjusted with sulphuric acid (Roth, Karlsruhe, Germany). The 96-well plates were incubated and growth was recorded using a Synergy H1 Hybrid Multi-Mode Reader (BioTek Instruments, Inc., Vermont, USA) for 30h at 37°C and 5% CO2.

### 8. Statistical Analysis

All experiments were performed with at least three independent biological replicates. To compare one variable between more than two groups, we performed One-way-ANOVA with Sidak (if not otherwise stated) as post-test to correct for multiple comparisons. Significant results are indicated with stars as: *p < 0.05; **p < 0.01; ***p < 0.001; ****p <0.0001. GraphPad Prism version 8 for Windows (GraphPad Software, San Diego, CA, USA, www.graphpad.com) was used for statistical analysis and visualization.

## Results

### 1. Multiple commensal bacteria isolated from pwCF inhibit the growth of *P. aeruginosa* lab strain PAO1

From CF sputum samples we isolated in total 83 aerobic or facultative anaerobic commensal bacterial strains, which comprise members that are generally not considered to be classical pathogens and recognized as part of the commensal airway microbiota according to literature ^29,30^ (sup. Table S1) as described previously ^25^.

A commensal isolate was identified as inhibitory if it prevented the growth of *P. aeruginosa* PAO1-mcherry throughout the 30h run time. Out of the screened CF commensal isolates, 35 isolates were identified as ‘inhibitory’ since they consistently prevented the growth of PAO1-mcherry during the 30h culture period (Fig. 1A). These inhibitory commensal isolates were all members of *Streptococcus* spp. In parallel to the measurement of RFU, we also assessed total bacterial growth via measurement of optical density at 600 nm (OD600). Representative results showing growth inhibition of PAO1-mcherry by *Streptococcus gordonii, S. mitis* and *S. cristatus* CF isolates in co-cultures are shown in Fig. 1B. For OD600 values, mono-culture of *P. aeruginosa* peaked above 2.0, while mono-culture of *Streptococcus* spp. did not exceed 0.6, making it also easy to assess growth inhibitory activities via OD600 value as shown in Fig. 1B.

**Fig. 1:**
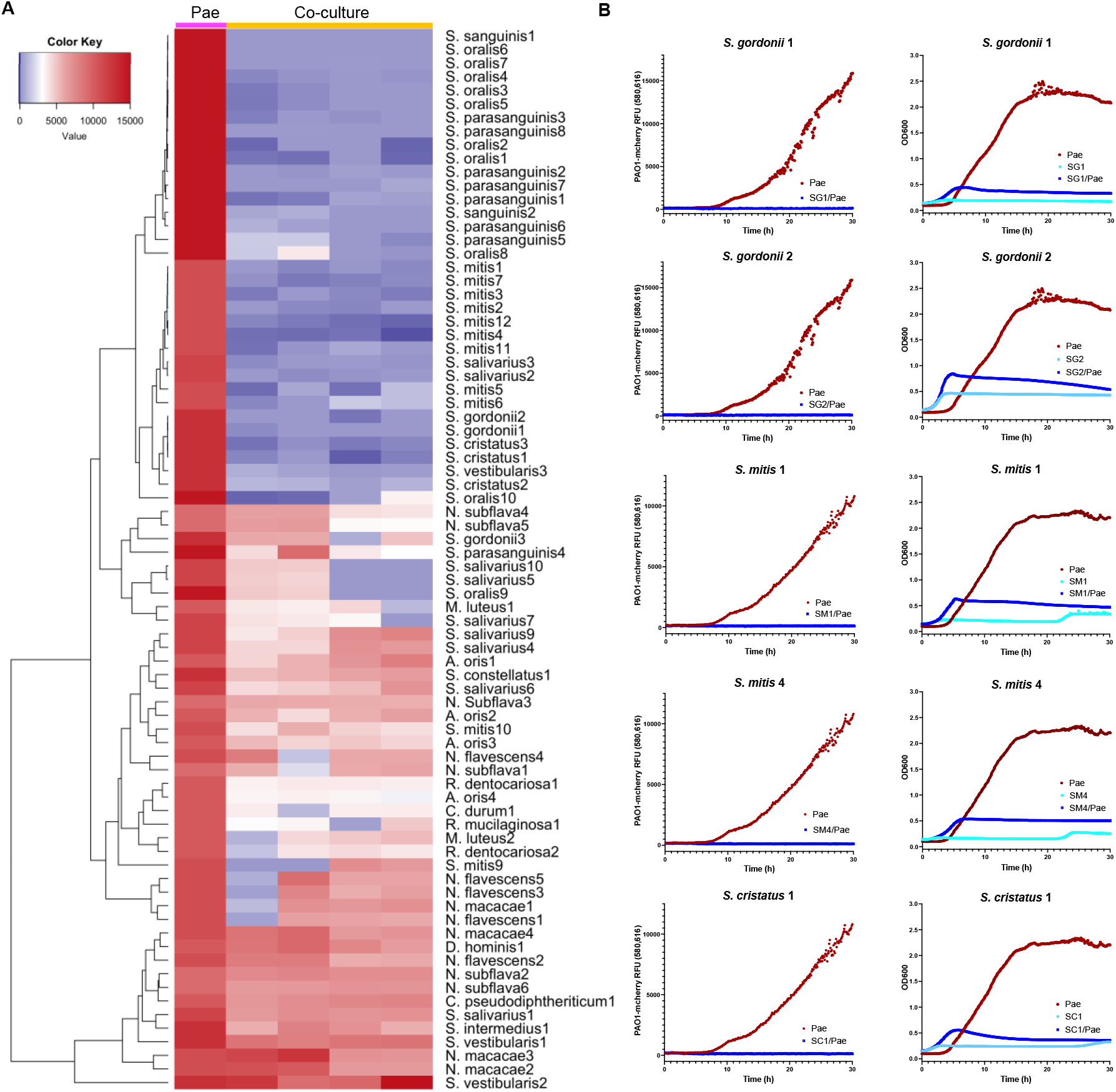
Various CF commensal strains inhibit the growth of *P. aeruginosa*. *P. aeruginosa* strain PAO1-mcherry (labelled as Pae) was mono-cultured or co-cultured with a commensal strain isolated from CF microbiome as indicated. Emission of red fluoresce (RFU) and OD600 were continually recorded for 30h. The inhibition is indicated by the low levels of RFU emission by PAO1-mcherry and low levels of OD600 in co-culture compared to that of *P. aeruginosa* mono-culture. **(A)** Heat map summary showing the RFUs after 30h growth of Psae in mono-or co-culture with S., *Streptococcus*; N., *Neisseria*; D., *Dermabacter;* A., *Actinomyces;* M., *Micrococcus*; R., *Rothia*; and C., *Corynebacterium*. Strains were numbered according to their isolation and identification, e.g., *S. mitis* 1 means *S. mitis* isolate1; n=4 biological replicates. **(B)** Representative results of growth curves from five *Streptococcus* isolates *S. gordonii* 1 (SG1), *S. gordonii 2* (SG2), *S. mitis* 1 (SM1), *S. mitis* 4 (SM4), and *S. cristatus* 1 (SC1). The left column shows the dynamic changes of RFU during the 30h culture period, and the right column shows the OD600 record during the 30h culture period; n=4 biological replicates.

Of note, some isolates belonging to *Rothia dentocariosa, R. mucilaginosa, Corynebacterium durum*, and *Actinomyces oris* also demonstrated intermediate antipseudomonal effects. However, unlike the inhibitory streptococcal isolates, they did not prevent the growth of PA01-mcherry throughout the 30h co-culture period and were therefore not identified as ‘inhibitory’. On the other hand, members belonging to the gram-negative *Neisseria* spp. demonstrated no potent antipseudomonal effects.

To identify isolates with the best antipseudomonal effects, the growth medium of the 30h co-cultures of commensal/PAO1 were plated on fresh CBA plates and incubated for an additional 24h to identify commensal isolates that could subsequently prevent the growth of PAO1-mcherry. No *P. aeruginosa* growth was detected in growth media after co-culture with the following eight strains: *S. mitis* 1, 3 & 4, *S. oralis* 2, 3 & 5 and *S. cristatus* 1 & 3.

### 2. Representative commensals inhibit the growth of several *P. aeruginosa* clinical isolates from pwCF at different disease progression stages

In addition to the common lab strain PAO1, which is a clinical isolate from wound ^49^, we also tested another commonly used lab strain *P. aeruginosa* ATCC27853, which is a clinical isolate from human blood sample ^50^. Importantly, multiple *P. aeruginosa* CF isolates from different disease progression stages were tested as well. Since the inhibitory effects on the growth of *P. aeruginosa* can be evaluated by OD600 values owing to the big difference in peak value between *P. aeruginosa* and commensals (as shown in Fig. 1B), OD600 read-out was used to demonstrate the inhibitory effects of commensals on the growth of *P. aeruginosa* clinical isolates (Fig. 2). The results of 4 representative commensal strains are displayed, among which *Streptococcus cristatus* 1 (SC1), *S*. mitis 1 (SM1) and *S. mitis* 4 (SM4) had strong inhibitory effects on the growth of *P. aeruginosa*, while *S. intermedius* 1 (SI1) had no such antipseudomonal effect and served as a control. Among the *P. aeruginosa* CF clinical isolates, PA24 is one non-mucoid early/intermittent CF isolate ^51^; PADD1 was isolated from one sample without detailed patient medical information; PA8, PADD2 and PADD3 are non-mucoid late/chronic CF isolates; CHA is a mucoid CF isolate ^52,53^. SC1, SM1 and SM4 displayed inhibitory effects on the clinical strain ATCC27853 and all the *P. aeruginosa* CF isolates from different disease progression stages, while SI1 did not inhibit any of these *P. aeruginosa* clinical isolates. Pseudomonal growth inhibition was also confirmed via plate count by inoculating the 30h co-cultured growth medium onto fresh CBA plates and further incubating for 24h. The results therefore demonstrate that selected streptococcal commensals have inhibitory effects on the growth of *P. aeruginosa* clinical isolates from pwCF at different disease progression stages.

**Fig. 2:**
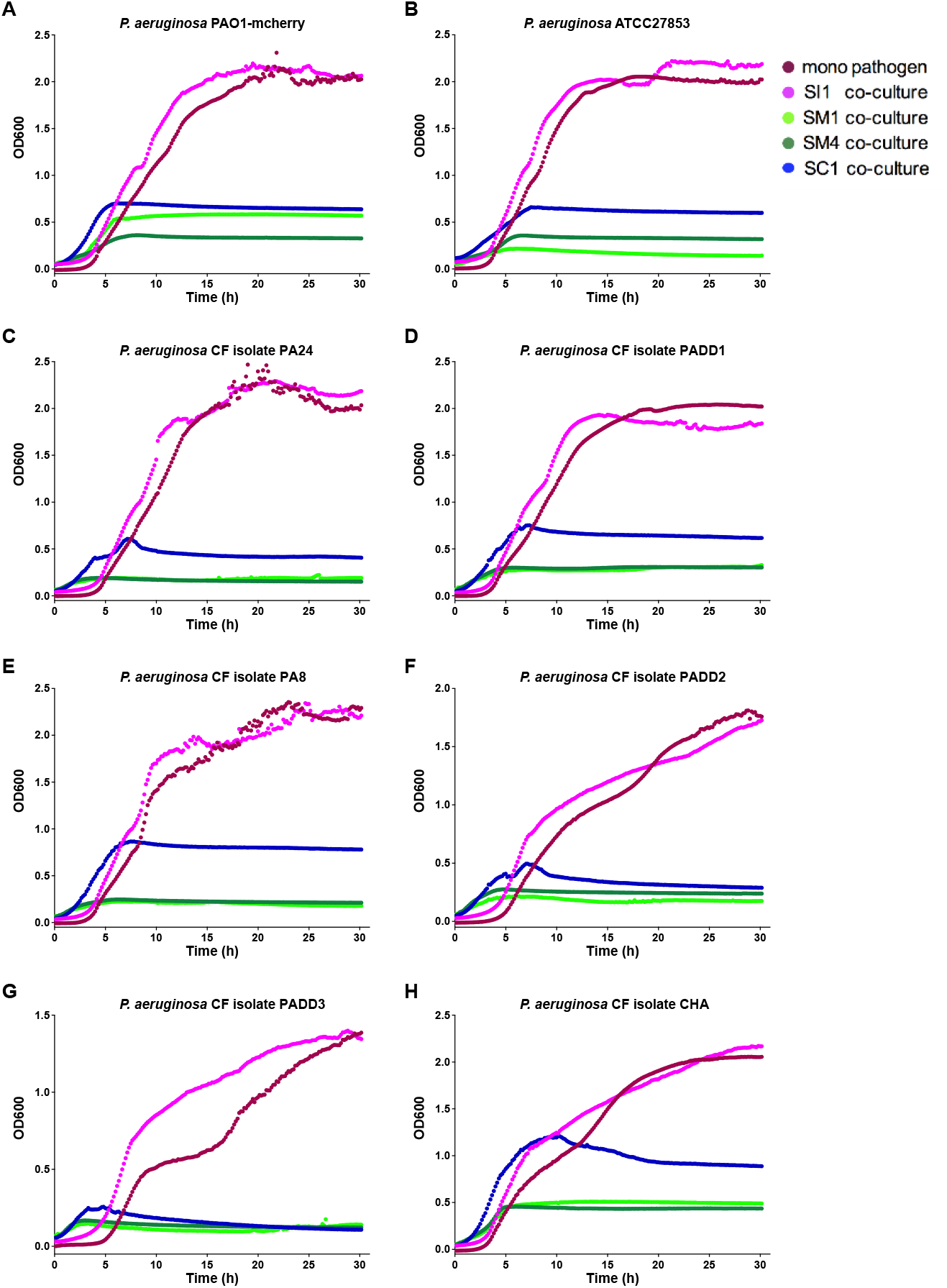
Representative streptococcal CF commensals inhibit the growth of different *P. aeruginosa* CF clinical isolates from different disease progression stages. Three commensal strains belonging to *S. mitis* (SM1, SM4) and *S. cristatus* (SC1) with growth inhibitory effects against PAO1-mcherry and one strain, *S. intermedius* (SI1), without inhibitory effects were tested for their effects on the growth of other *P. aeruginosa* CF isolates. **(A&B)** Two *P. aeruginosa* non-CF clinical isolates PAO1 (wound isolate) and ATCC27853 (blood isolate). **(C)** PA24: non-mucoid early/intermittent CF isolate. **(D)** PADD1: the patient condition is unclear. **(E, F&G)** PA8, PADD2 and PADD3: non-mucoid late/chronic CF isolates. **(H)** CHA: mucoid CF isolate.

### 3. Representative commensals inhibit the growth of other non-pseudomonal CF pathogens

In addition to *P. aeruginosa*, multiple other pathogens are also frequently detected in the CF airway infections ^54^, including gram-negative pathogens like *Klebsiella pneumoniae*, *Haemophilus influenzae*, *Proteus mirabilis*, *Achromobacter xylosoxidans*, *Burkholderia multivorans*, and *Stenotrophomonas maltophilia* as well as gram-positive pathogens like *Staphylococcus aureus* and *Enterococcus faecalis*. Therefore, selected commensals (SM1, SM4 & SC1) were co-cultured with clinical isolates of other non-pseudomonal CF pathogens to analyze if growth inhibitory effects could also be observed, while the non-inhibitory SI1 served as a control.

With a similar setting as described in results section 2 using OD600 read-out, we first studied several widely-used lab strains: *Klebsiella pneumoniae* ATCC13883, *Haemophilus influenzae* ATCC49247, *Enterococcus faecalis* ATCC29252, *Staphylococcus aureus* ATCC25923, and observed growth inhibition of these pathogens (the results are shown in sup. Fig. S1). We then evaluated multiple CF clinical isolates, namely *S. aureus* SPA4, *E. faecalis* ETF1, *K. pneumoniae* KP1, *P. mirabilis* PTM1, *Achromobacter xylosoxidans* AX1, *Burkholderia multivorans* BM1, and *Stenotrophomonas maltophilia* STM2, and also observed growth inhibition mediated by SM1, SM4 & SC1 (Fig. 3A-G). Taking the results together, it was observed that the inhibitory activity mediated by selected CF streptococcal commensals is not limited to *Pseudomonas* spp. but occurs across a wide range of typical CF pathogens.

**Fig. 3:**
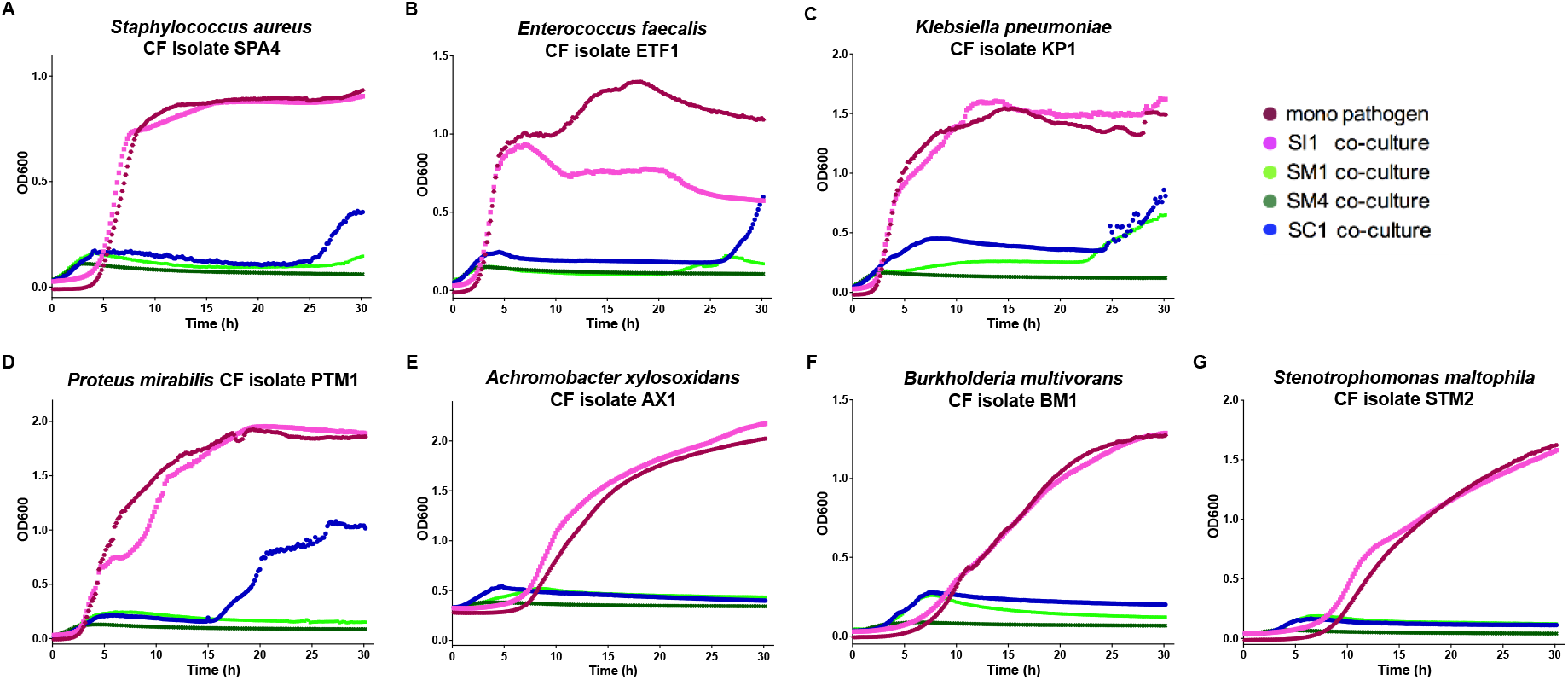
CF commensals inhibit the growth of various other CF pathogens. Representative results from three inhibitory commensal strains SM1, SM4 and SC1 and one non-inhibitory commensal strain SI1 are displayed. (**A-G**) Multiple CF isolates of typical pathogens were co-cultured with CF commensals, and the inhibitory effects are displayed as low levels of OD600 in co-culture compared with that of pathogen mono-culture. The strains tested are *Staphylococcus aureus* SPA4 (A), *Enterococcus faecalis* ETF1 (B), *Klebsiella pneumoniae* KP1 (C), *Proteus mirabilis* PTM1 (D), *Achromobacter xylosoxidans* AX1 (E), *Burkholderia multivorans* BM1 (F), *Stenotrophomonas maltophilia* STM2 (G).

### 4. Commensal-conditioned medium inhibits pathogen growth with pH playing a role, but not iron competition or peroxide production, and the inhibitory effect is not heat sensitive

To investigate mechanisms by which commensals interfere with *P. aeruginosa*, we performed experiments with commensal-conditioned medium (CM) which contained molecules secreted by commensal bacteria into the growth medium. The results showed that the growth-inhibitory effects could be observed when using CM, and the inhibitory effects were not affected by heat treatment (90°C for 30 minutes) (sup. Fig. S2). This indicates that the inhibitory commensals could release certain active compounds into their growth media which can mediate inhibitory effects, yet these compounds are not heat-sensitive. Furthermore, the CMs were also pretreated overnight at 56°C with 100 μg/ml proteinase K, which would cleave peptide bonds and thereby disintegrate protein molecules. The proteinase K-treated CM still preserved the growth inhibitory effects (sup. Fig. S2) indicating that the released inhibitory compound is not proteinaceous.

Iron is required by *P. aeruginosa* for several cellular pathways. *Streptococcus* spp. have developed mechanisms to collect iron from their environment ^55^. It was therefore analyzed if the observed antipseudomonal effect mediated by streptococcal commensals was due to iron sequestration which could make iron unavailable for pseudomonal growth. Varying concentrations (5 – 1,000 μM) of ammonium ferrous sulfate (AFS) were first added to the growth medium of PAO1-mcherry to ensure the concentrations were not toxic. After confirming these iron concentrations did not alter the growth of PAO1-mcherry, they were added to the CM of SM4. Irrespective of the added iron concentration (-excess or limited-) the SM4 CM was found to prevent the growth of PAO1-mcherry after incubation for 30h (sup. Fig. S3).

Since peroxide production is one of the possible mechanisms proposed in the literature by which members of *Streptococcus* spp. mediate growth inhibitory effects, the peroxide concentrations of the CMs from the inhibitory SM1, SM3 & SM4 were analyzed (sup. Fig. S3). A concentration of 0.2 mg/ml catalase was then added to the CMs and their peroxide concentrations were confirmed to be eliminated. However, these catalase-treated CMs still prevented the growth of PAO1-mcherry for 30 h (sup. Fig. S3) demonstrating that peroxide production is not a major mechanism.

Moreover, we measured pH values of CMs of multiple commensal CF isolates with or without growth-inhibitory effects, and the results (sup. Table S4) revealed that all the strains with growth-inhibitory effects had a low pH value (range 4.78-5.28), whereas strains without growth-inhibitory effects usually had a higher (neutral) pH value. When the pH of these inhibitory CMs was adjusted to 7.0, they all lost their growth inhibitory effects (sup. Fig. S3). However, a low pH of 4.5 itself was not sufficient to prevent the growth of PAO1-mcherry (sup. Fig. S3), suggesting a low pH value might be a necessary supporting condition for the inhibitory effects.

### 5. Comparative genomic analysis reveals functions highly enriched in bacteria that could inhibit the growth of CF pathogens

The results of the screening study (as shown in Fig. 1) indicate the inhibitory effects are strain-specific. Therefore, we explored the differences between strains with and without inhibitory effects. Since we had obtained a profound *Streptococcus* CF-isolate collection and also observed differences among the *Streptococcus* strains, it became possible to investigate genetic differences between inhibitory and non-inhibitory strains in a shared genetic background. For that, we selected 9 strains without inhibitory effects (*S. vestibularis* 1 (SV1), *S. vestibularis* 2 (SV2), *S. salivarius* 4 (SS4), *S. salivarius* 6 (SS6), *S. salivarius* 9 (SS9), *S. parasanguinis* 4 (SP4), *S. mitis* 10 (SM10), *S. intermedius* 1 (SI1), and *S. constellatus* 1 (SL1)), defined as “negative group”; and 9 strains with strong inhibitory effects (*S. cristatus* 1 (SC1), *S. gordonii* 1 (SG1), *S. gordonii* 2 (SG2), *S. oralis* 2 (SO2), *S. oralis* 4 (SO4), *S. mitis* 1 (SM1), *S. mitis* 4 (SM4), *S. parasanguinis* 1 (SP1), and *S. parasanguinis* 8 (SP8)), defined as “positive group” to perform comparative genomic analysis. These strains were sequenced to acquire their complete whole genome sequences. The strain taxonomy was further investigated through comparing the sequences of these strains with publicly available whole genome sequences from type strains belonging to the genus *Streptococcus* (sup. Fig. S4, with details described in the supplementary material), which indicates the correct taxonomy of SM10 and SL1 are *S. pseudopneumoniae* and *S. sanguinus*, respectively. In the following analyses, we replaced the name of SM10 with SPS1(*S. pseudopneumoniae* 1) and replaced the name of SL1 with SN3 (*S. sanguinis* 3).

We then compared the whole genome sequences of these strains to search for functions that might be associated with the inhibitory effects by identifying functions that are highly enriched in the positive group, while largely missing in the negative group (based on function annotation from the NCBI COG database COG20). For that, we conducted pan-genome analysis of the 18 strains of the positive group and the negative group (Fig. 4A) and carried out comparative functional enrichment analysis. The results of these analyses revealed functions highly enriched in the strains with inhibitory effects (Fig. 4B, with details listed in sup. Table S6), and also revealed functions highly enriched in the strains without inhibitory effects (sup. Fig. S5, with details listed in sup. Table S7). Top five enriched function categories in each group are displayed according to their relative ratio (based on the number of functions in each category). A clear difference was detected between the positive group and the negative group. In the positive group, the most highly enriched function category is “carbohydrate transport and metabolism”, whereas in the negative group, the most enriched function category is “defense mechanisms”. The details of functions enriched in the category “carbohydrate transport and metabolism” with pathway identifications are listed in Fig. 4C. Regarding signaling pathways, “Pentose phosphate pathway” was highly enriched in the strains with inhibitory effects, and “Gluconeogenesis” is also enriched. These results provide insights that the inhibitory effects might be mainly derived from the ability to produce or utilize carbohydrate sources.

**Fig. 4:**
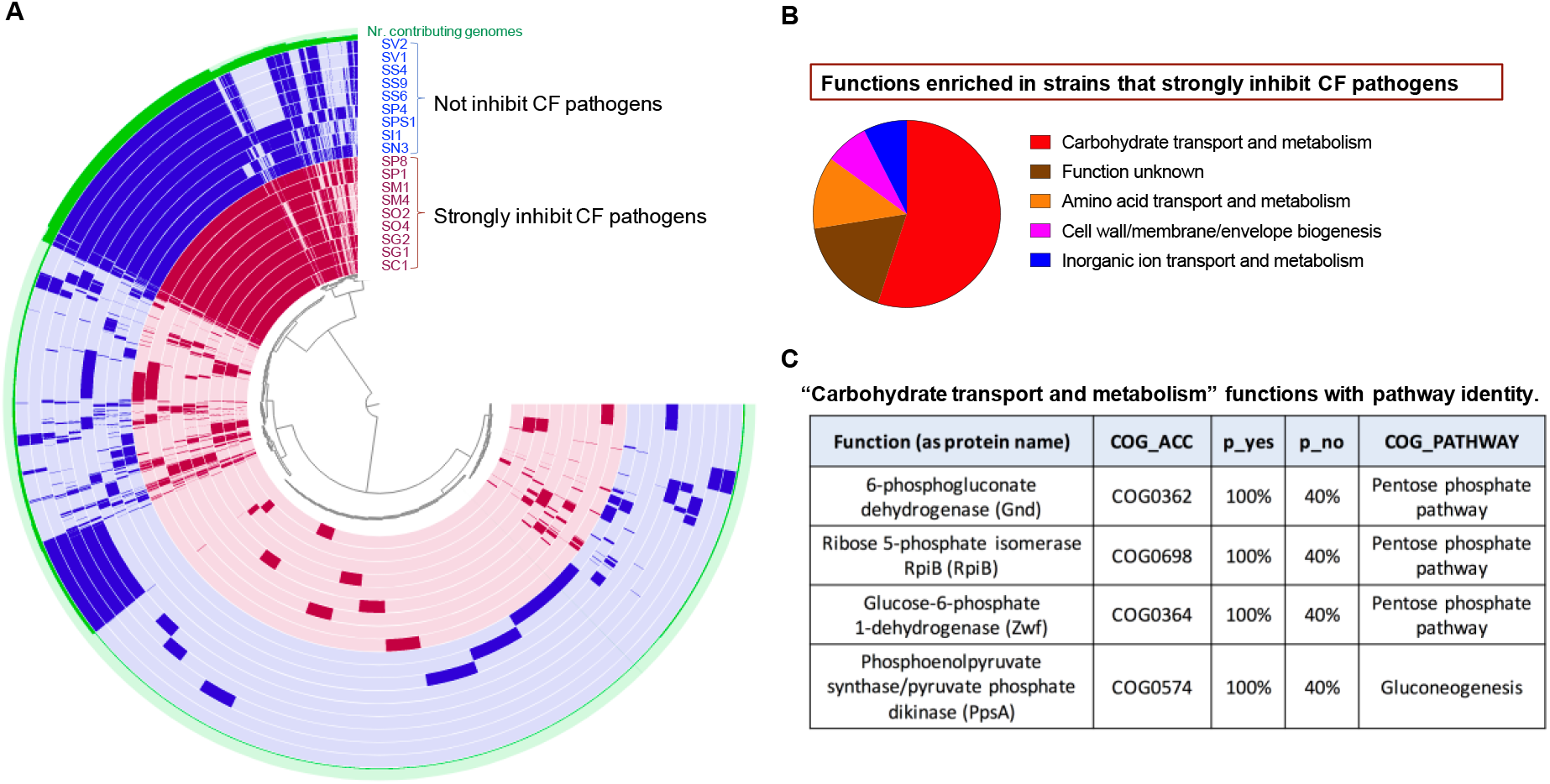
Comparative genomic analysis of *Streptococcus* strains reveals functions highly enriched in strains with inhibitory effects on the growth of *P. aeruginosa*. A. Pan-genome analysis to compare strains with inhibitory effects (the positive group) and strains without inhibitory effects (the negative group). One layer represents one genome, with one extra layer showing number of contributing genomes to each orthologous gene family (labelled as “Nr. contributing genomes”). Genomes of strains showing inhibitory effects (SP8, SP1, SM1, SM4, SO2, SO4, SG2, SG1, SC1) are colored in red, and genomes of strains without inhibitory effects (SV2, SV1, SS4, SS9, SS6, SP4, SPS1, SI1, SN3) are colored in blue. The clustering is based on presence/absence of orthologous gene family using Euclidean distance and Ward Linkage. B. Top five function categories highly enriched in the positive group. The relative ratios of the number of functions in each category are displayed as a pie chart. C. Functions enriched in the positive group in the category “carbohydrate transport and metabolism” with pathway identifications. p_yes and p_no indicate the frequency of the relevant function being detected either in the “positive group” or in the “negative group”. COG_ACC is the accession number in the COG database, while COG_PATHWAY is the pathway in the COG database the relevant function is classified in.

### 6. Commensals mediate growth-inhibition effects via the production of a short-chain fatty acid, acetate

In the context of CF airway, the major carbon source for *P. aeruginosa* is phosphatidylcholine (an airway surfactant) and the primary breakdown products of phosphatidylcholine are glycerol and acetate, the amount of which in the microenvironment can regulate *P. aeruginosa* metabolism ^56,57^. In particular, it has been shown that an acetate-rich environment may attenuate the “EDEMP cycle” including the Entner-Doudoroff pathway, the Embden-Meyerhof-Parnas pathway, and the pentose phosphate pathway in *P. aeruginosa* ^56^. Acetate is one sort of short-chain fatty acids (SCFAs). Previous studies indicated that SCFAs produced by commensals or from other sources may interfere with the growth of pathogens ^58–61^.

As revealed by comparative genomic analysis (Fig. 4C), the “pentose phosphate pathway” was highly enriched in commensal strains with inhibitory effects, the activation of which may lead to high production of acetate or a few other SCFAs ^62^. This raised the hypothesis that the activation of pentose signalling pathways in certain commensals in CF airways leads to higher production of acetate, which (when exported to the microenvironment) may switch off the “EDEMP cycle” in *P. aeruginosa* and consequently affect growth.

We analysed if commensals with growth inhibitory effects may produce SCFAs. We selected two CF commensal strains that consistently displayed strong inhibitory effects (SM3 and SM4) as well as one strain with an intermediate inhibitory effect (RM1). As shown in Figure 5, in the conditioned medium (CM) of SM3 and SM4, the amount of acetate was two to three times higher than that in the culture medium Columbia Broth (CB) (uninoculated). On the other hand, the acetate amount in the CM of *P. aeruginosa* was not higher than that in the culture medium. The amount of acetate in the CM of RM1 was lower than that of SM3 and SM4, but also much higher than that of *P. aeruginosa* and the culture medium. For other SCFAs measured, including 2-methyl-butyrate, i-valerate, caproate, propionate, butyrate, i-butyrate and valerate, no clear differences were detected between CM of commensals and the culture medium. Taken together, the analyses of SCFAs in commensal-CM revealed that SM3, SM4 and RM1 may release a large amount of acetate to the microenvironment (range: 1.2-6.0 mM), which might impact other bacteria living in the same microenvironment.

**Fig. 5:**
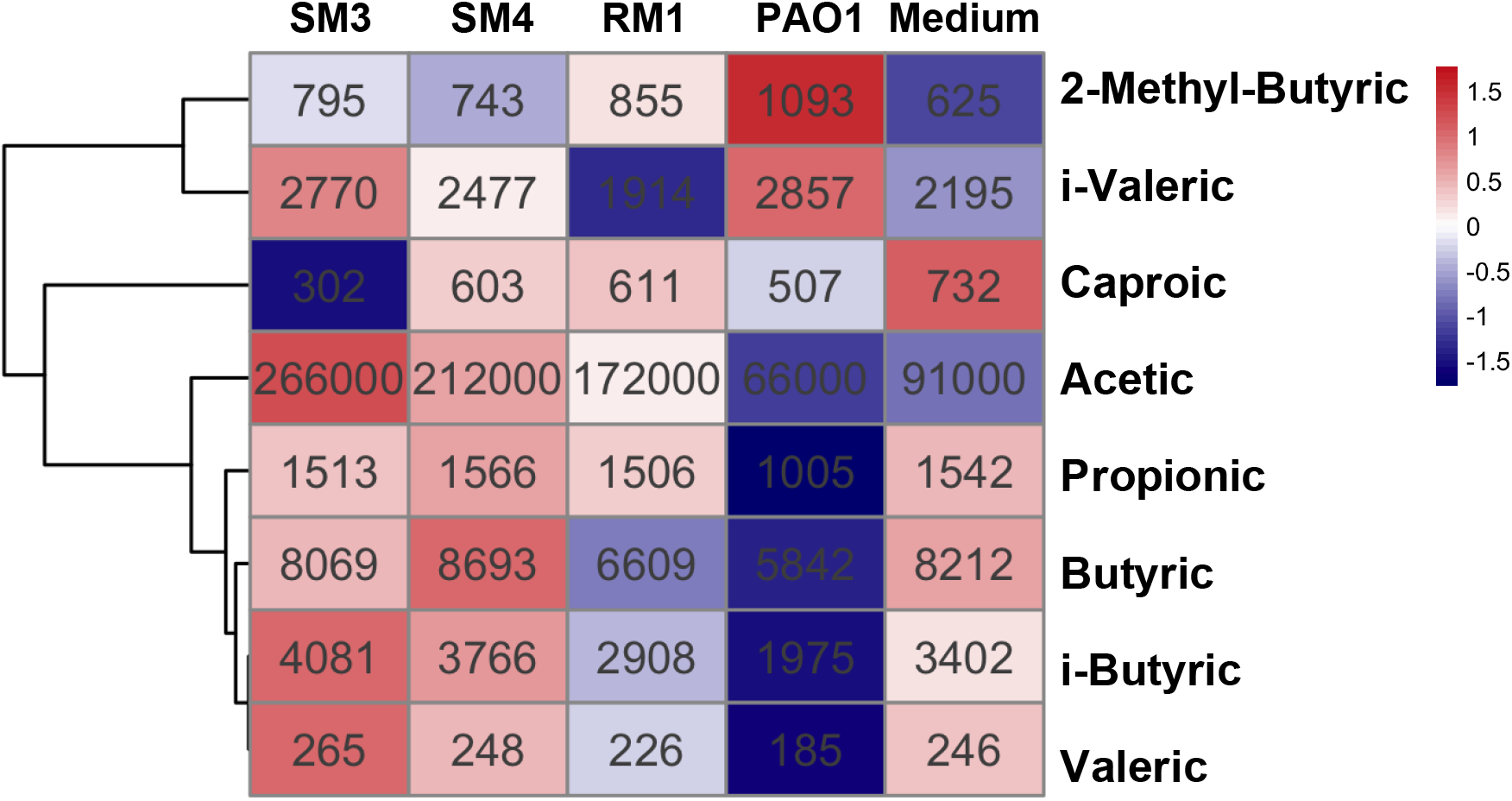
Profiling of SCFAs in conditioned media of *P. aeruginosa* and commensals with inhibitory effects. The heatmap displays Z-score transferred values, while the original concentrations are labelled in each plot with unit **ng/ml**. “Medium” is the culture medium Columbia Broth (CB) without any bacteria. PAO1, RM1, SM3, SM4 refer to the conditioned media of them.

### 7. Exogenous addition of acetate at pH 5.0 inhibits the growth of *P. aeruginosa*

Previous studies indicated that during growth on the different carbon sources, a global remodeling can take place in *P. aeruginosa* and thereby many aspects of the bacterial behavior could be affected ^56^. To confirm that acetate indeed plays a role in the observed antipseudomonal effect, PAO1-mcherry was grown in the presence of added acetate with concentrations close to that released by commensal-CM (1.2–6.0 mM). The pH was also adjusted to 5.0 to mimic the pH of commensal-CM with inhibitory effects. The adjustment of pH itself did not result in inhibitory effects: at a neutral pH, PAO1-mcherry alone has a lag phase of 3.4 h in the culture medium CB, and at a pH 5.0, the lag phase only slightly extended to 5.6 h in CB (Fig. 6A). With 1.2 mM acetate in CB at pH 5.0, the lag phase was increased to 7.5 h, while the 6.0 mM concentration drastically increased this to 24.3 h. Multiple *P. aeruginosa* CF isolates were also tested using 6.0 mM acetate in CB at pH 5.0. Under this condition, the CHA, PA8, PA24, PADD1 and PADD2 isolates were inhibited for 24.4, 26.1, 23, 22 and >30 h respectively (Fig. 6B-D). On the other hand, growing PAO1-mcherry in CB at a normal neutral pH with a higher concentration of 17 mM acetate had no clear impact on the growth of PAO1 (data not shown). The results indicate acetate may inhibit the growth of pathogens in a low-pH dependent manner.

**Fig 6:**
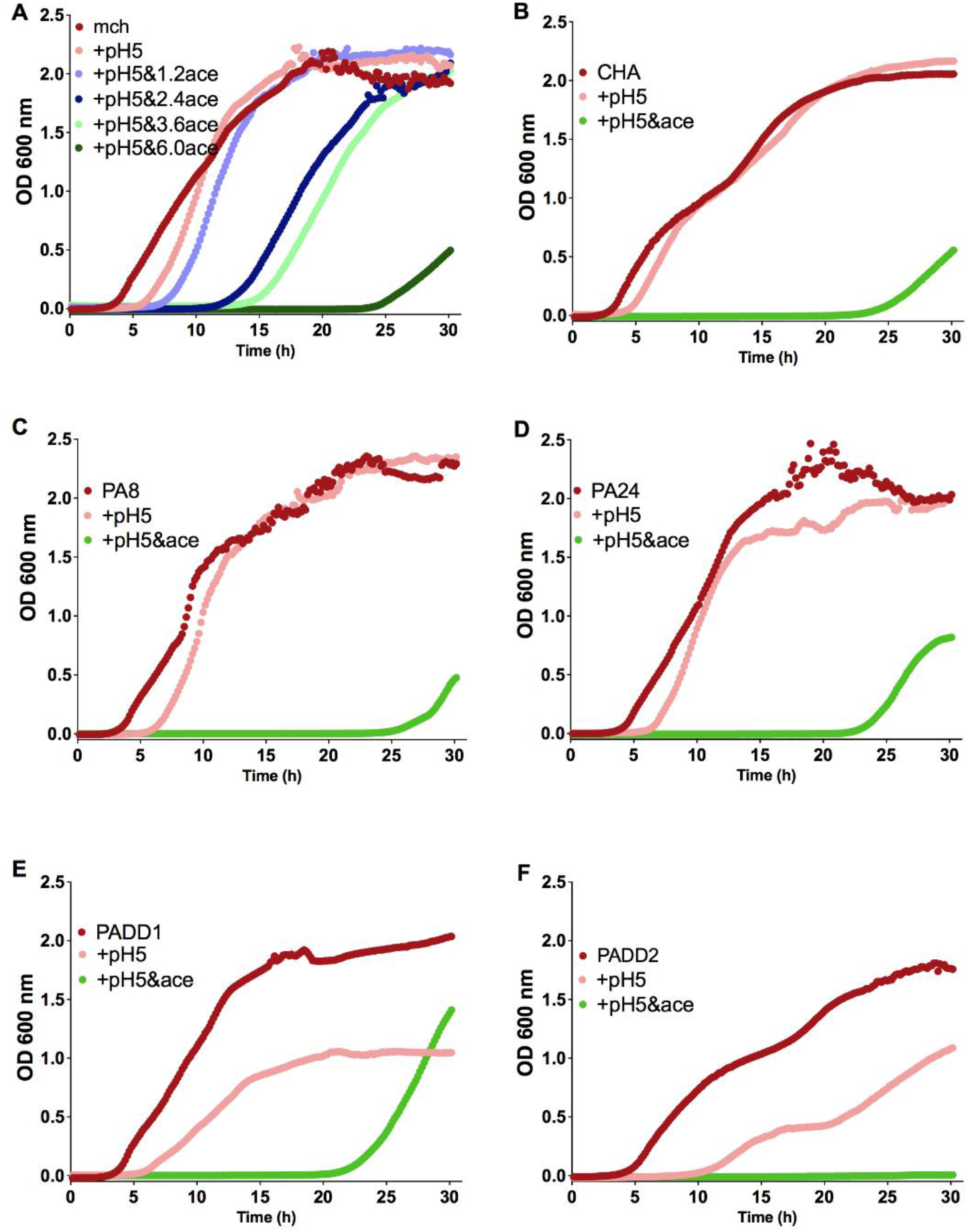
The impact of SCFA acetate on the growth of *P. aeruginosa*. **A.** Different concentrations of acetate were added to the culture media CB and the growth of *P. aeruginosa* PAO1-mcherry was displayed with the readouts of OD600. **B-F.** The impact of 6.0 mM acetate addition in CB on the growth of *P. aeruginosa* CF isolates CHA (B), PA8 (C), PA24 (D), PADD1 (E) and PADD2 (F).

## Discussion

Interactions between pathogens and commensals in CF airway infections are of increasing interest. Although multiple studies suggest that higher microbial diversity in the airway is in favor of lung function preservation, current commonly applied antibiotic treatment targeting certain pathogen infection often largely diminishes microbial diversity in the airway. Deciphering the nature of the interaction between pathogens and commensals in the airway could be used to guide CF infection treatment strategies or develop novel microbiome-based therapies. In this study, we discovered that various CF airway commensal strains strongly inhibited the growth of *P. aeruginosa* and multiple other CF pathogens. The SCFA acetate, a metabolite released by commensals, played a critical role in mediating the inhibitory effects on pathogen growth.

The results of this study concur with previous investigations: in a screening study from another group, bacterial strains were isolated from the microbiota of posterior pharyngeal walls of healthy subjects^63^, among which four inhibitory isolates against common respiratory pathogens were identified: two from *S. mitis* and two from *S. parasanguinis*. Here we observed notable inter- and intra-species variations in the streptococcal-mediated growth inhibitory effects: members belonging to the Mitis (*S. mitis, S. oralis* and *S. parasanguinis*) and Sanguinis (*S. sanguinis, S. gordonii*, and *S. cristatus*) groups were found to be generally more inhibitory than members belonging to the Salivarius (*S. salivarius* and *S. vestibularis*) and Anginosus group (*S. intermedius* and *S. constellatus*) group. Consistent with our results, members of the Mitis and Sanguinis group were shown in one previous study to inhibit the growth of *P. aeruginosa*^64^. In that study *S. parasanguinis* was also demonstrated to protect *Drosophila melanogaster* from death caused by *P. aeruginosa* infection ^64^. Members of the Anginosus group, however, have been implicated in CF pulmonary exacerbations and shown to increase the pathogenicity of *P. aeruginosa* by upregulating the expression of virulence gene factors ^22,65^. *S. salivarius* and *P. aeruginosa* may form dual biofilms via the presence of a maltose-binding surface protein (MalE) ^66^. Intriguingly, there are *Salivarius* strains like *S. salivarius* strain (M18) which has been demonstrated to inhibit *P. aeruginosa* ^67^. This provides an example of the dynamic and complex commensal-pathogen and strain-specific relationships that exist in the airways.

To understand the underlying mechanisms behind the growth-inhibition effects of commensals, CMs from selected inhibitory commensals were obtained and were found to inhibit the growth of several clinical *P. aeruginosa* isolates. This observation indicates that these bacteria can condition their environment to prevent the growth of pathogens. Other studies have also reported growth-inhibitory effects mediated by CMs from different commensals but could not yet identify the responsible molecules ^67–69^. Our results indicate that commensals may mediate growth-inhibitory effects by producing acetate and lowering pH. The most common SCFAs, acetic, propionic, and butyric acid, are weak acids with pKa values of 4.76, 4.87, and 4.81 respectively ^70^. In the undissociated state, weak acids like SCFAs have been shown to exhibit increased antimicrobial effects at low pH ^71^. The most common explanations for the antimicrobial activities are the uncoupling effect and the consequent disruption of anion pools^71–74^, which were suggested to affect metabolism and growth of the bacteria.

The production of bacteriocins or reactive oxidative radicals like H2O2 and nitrite is frequently proposed in literature as the major antagonistic tools employed by oral streptococci against pathogens ^23,75^. Our results indicate that the production of SCFA acetate is one of the major mechanisms employed by *Streptococcus* spp. and other commensals like *Rothia mucilaginosa* in mediating growth-inhibition effects. SCFAs, the major ones being acetate, propionate, and butyrate, are usually described as the primary metabolites produced by the gut microbiota from anaerobic fermentation of indigestible polysaccharides ^76–78^. Although the protective role of SCFAs in the gut has been well established, their role in respiratory infections is only emerging and still controversial as some studies argue for ^79^ and a few others against ^80^ a protective effect. Some studies proposed that the gut-lung axis is a possible route via which SCFAs produced in the gut migrate into the airways to mediate protective effects ^58,76,81^. In the present study, it is demonstrated that members of the lung microbiota also produce SCFAs like acetate, indicating that commensal members of the lung microbiota may contribute to the lung SCFAs concentration.

It has been shown *in vitro* that *S. mitis, S. oralis, S. gordonii*, and *S. sanguinis* inhibited the growth of *P. aeruginosa* only when they were introduced as primary colonizers but when introduced as secondary colonizers after the introduction of *P. aeruginosa* as the primary colonizer, they failed to inhibit *P. aeruginosa* growth ^23^. Considering the results of this study, the reason for this phenomenon might be that commensals are unable to successfully grow and secrete metabolites like acetate to mediate protective effects if they are secondary colonizers. Since the antagonistic effect of acetate is pH dependent, the use of acidifying compounds together with a therapeutic acetate dosage could hold potential in enabling the decolonization of CF pathogens and recolonization by protective commensals.

More studies are needed to compare the SCFA contents in CF and healthy patients and to investigate how the airway SCFAs composition is altered during pulmonary exacerbations. Also, SCFA levels might be linked to the microbiome structure. Such studies will decipher the roles that individual SCFAs play in CF health and disease. Furthermore, since the formation of biofilms is a common way by which *P. aeruginosa* promotes its chronic presence in the airways, the effect of acetate on pseudomonal biofilms should be the topic of further studies.

In summary, this study demonstrates that various airway commensal bacteria may inhibit the growth of pathogens in the CF airway via the production of acetate which can antagonise invading pathogens via a pH-dependent mechanism. These findings indicate that commensal bacteria in the airway of pwCF could help control the growth of pathogens and highlight the importance of preserving airway commensals when designing infection treatment strategies. In particular, attention should be paid when choosing antibiotics to minimize side effects on beneficial commensal bacteria.

## Acknowledgments

We thank Dr. Anne Blanc-potard from the Centre National de la Recherche Scientifique (CNRS) for kindly providing us the *P. aeruginosa* strain PAO1 pBPF-mcherry. We thank Selina Mayer for the technical support.

