## Supplementary material for "Commensal bacteria can inhibit the growth of *P. aeruginosa* in cystic fibrosis airway infections through a released metabolite": sup.

#### 1. Results

##### 1.1 Taxonomy investigation of *Streptococcus* CF isolates

We compared the sequences of the strains with publicly available whole genome sequences from type strains belonging to the genus *Streptococcus* (sequence information of the type strains is listed in Suppl. Table S5) with pairwise average nucleotide identity (ANI) analysis (Fig. S4A). This index is a robust similarity metric that has been broadly used to delineate inter- and intra-species strain relatedness (Jain, Rodriguez, Phillippy, Konstantinidis, & Aluru, 2018). When compared with the genomes of the type strains, most of these *Streptococcus* CF isolates showed the highest ANI value when paired with the type strain of the species identified through MALDI-TOF, with two exceptions: SM10 and SL1. SM10 showed the highest ANI value when paired with the type strain of *S. pseudopneumoniae,* while SL1 showed the highest ANI value when paired with the type strain of *S. sanguinis*.

We further performed phylogenomic analysis of these *Streptococcus* genomes based on core-genome sequence analysis (Fig. S4B). Similarly, most CF strains were closely clustered together with the type strains from the same species as being identified through MALDI-TOF, except for SM10 and SL1, which clustered with the type strains of *S. pseudopneumoniae* and *S. sanguinis,* respectively. This means the results of both ANI analysis and phylogenomic analysis consistently support the correct taxonomy of SM10 and SL1 as *S. pseudopneumoniae* and *S. sanguinus*. In the following analyses, we replaced the name of SM10 with SPS1(*S. pseudopneumoniae* 1) and replaced the name of SL1 with SN3 (*S. sanguinis* 3).

Streptococci have been classified into several groups based on their phylogenomic relationship (Richards et al., 2014). Based on previously identified classification of type strains, from both phylogenomic analysis and ANI analysis, *Streptococcus* CF isolates were classified into specific groups based on their clustering with specific type strain. Combining the information from Fig. 1 and Fig. S4, it appears that in our collection, *Streptococcus* CF isolates were mainly from the Salivarius, Mitis, Anginosus and Sanguinis groups, and rarely from the Bovis or Pyogenic group. The combined data show that members belonging to the Mitis (*S. mitis, S. oralis* and *S. parasanguinis*) and Sanguinis (*S. sanguinis, S. gordonii,* and *S. cristatus*) group demonstrate more growth inhibiting effects against CF pathogens than members belonging to the Salivarius (*S. salivarius* and *S. vestibularis*) and Anginosus (*S. intermedius* and *S. constellatus*) group.

### 2 Supplementary Figures and Tables

#### 2.1 Supplementary Figures

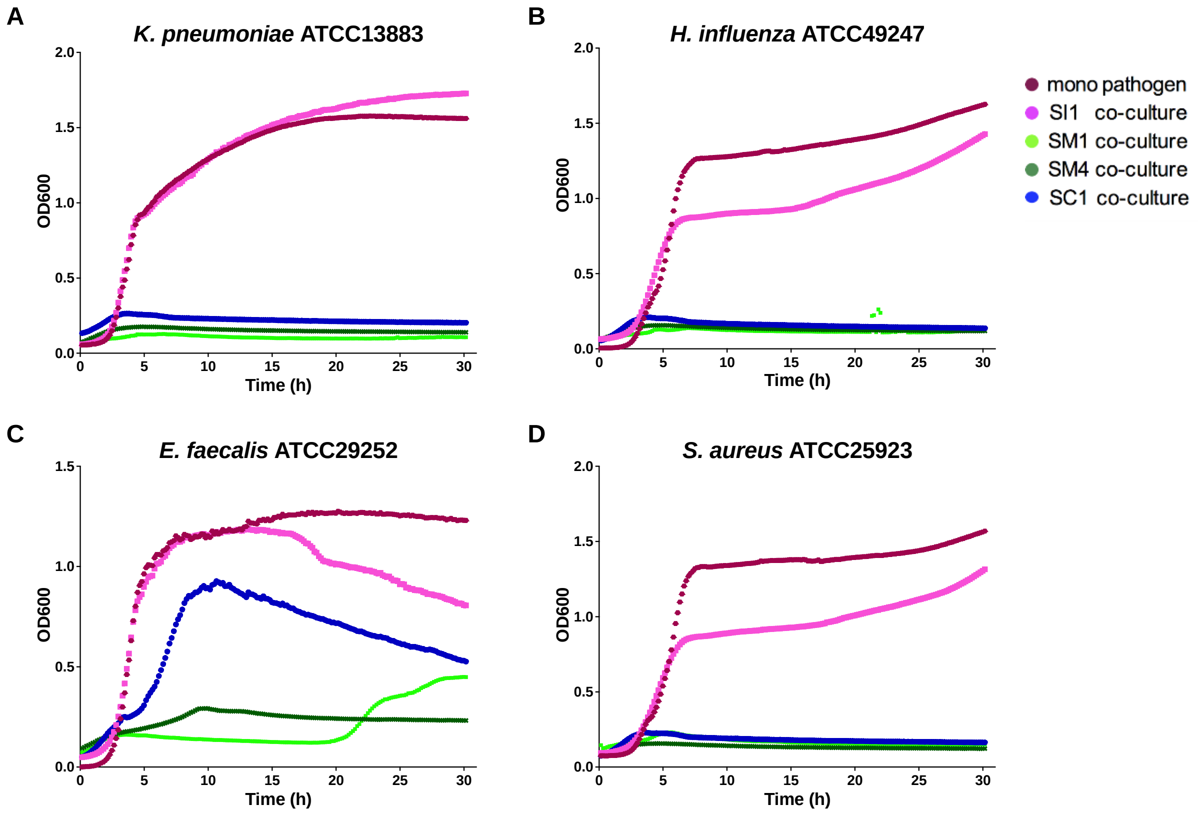


**Figure S1**: **CF commensals could inhibit the growth of various typical CF pathogens.** Representative results from three positive commensal strains SM1, SM4 and SC1 and one negative commensal strain SI1 are displayed. (**A-D**) several typical CF pathogens were co-cultured with CF commensals, and the inhibitory effects are displayed as low levels of OD600 in co-culture compared with that of pathogen mono-culture. The strains tested are widely used lab strains *Klebsiella pneumoniae* ATCC13883 (A), *Haemophilus influenzae* ATCC49247 (B), *Enterococcus faecalis* ATCC29252 (C), *Staphylococcus aureus* ATCC25923 (D).

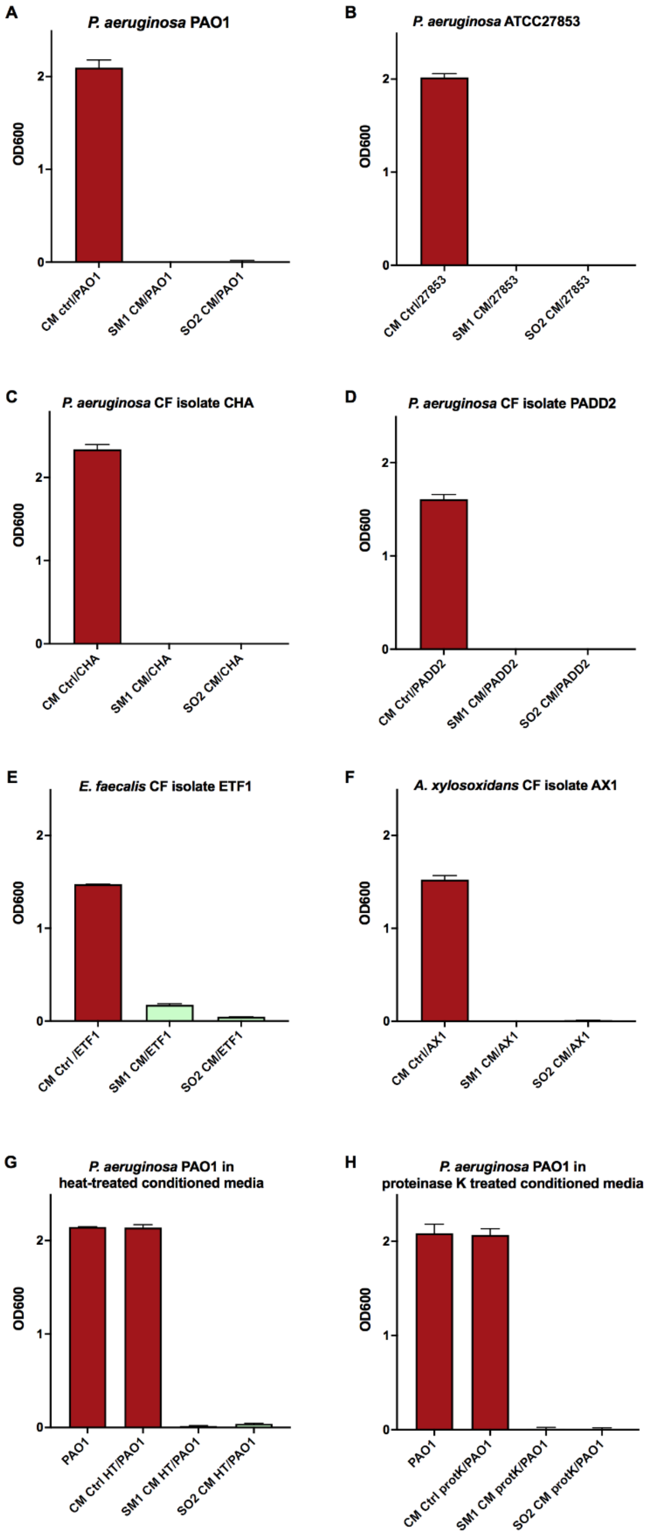


**Figure S2: Conditioned media of commensals with growth-inhibitory effects could also inhibit pathogen growth, and the inhibitory effects were neither affected by heat treatment (90 °C for 30 minutes) nor affected by the treatment of Proteinase K (protK).** Conditioned media of SM1 and SO2 could inhibit the growth of *P. aeruginosa* non-CF strains PAO1 (A) and ATCC27853 (B), as well as CF strains CHA (C) and PADD2 (D), and other typical CF pathogens, e.g. *E. faecalis* CF isolate ETF1 (E) and *A. xylosoxidans* CF isolate AX1 (F). The inhibitory effects were not affected by heat treatment (90 °C for 30 minutes) (G), indicating the effect was not heat sensitive. Also the treatment of 100 μg/ml proteinase K overnight at 56 °C could not affect the inhibitory effects (H), indicating the effect was not proteinaceous.

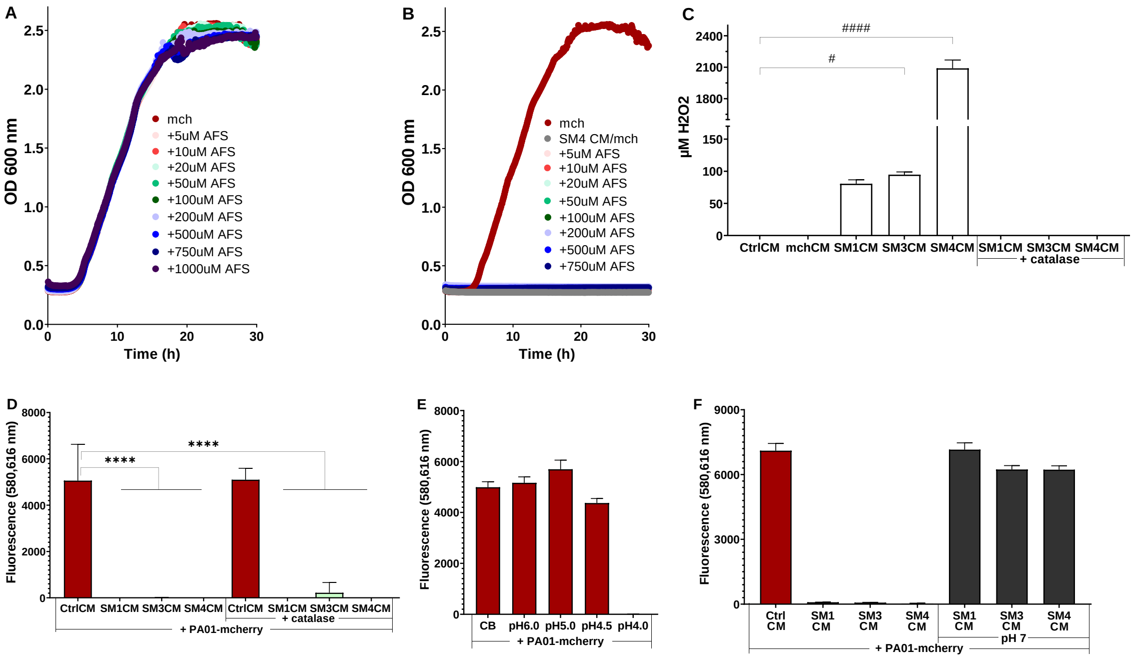


**Figure S3: pH plays a role but not iron competition or peroxide production in the commensal CM-mediated *P. aeruginosa* growth inhibition.** 5, 10, 20, 50, 100, 200, 500, 750, and 1,000 µM of ammonium ferrous sulfate (AFS) was added to (A) Columbia broth (CB) as control and (B) *S. mitis* 4 conditioned medium (SM4 CM), and PA01-mcherry (mch) was cultured in these iron-fortified media. (C) $H_2O_2$ concentration of the selected *S. mitis* CMs (SM1 CM, SM3 CM & SM4 CM) either alone or treated with 0.2 mg/ml catalase. Fluorescence reads at 30 h showing mch growth in (D) selected *S. mitis* CMs treated with or without 0.2 mg/ml catalase, (E) CB at neutral pH or adjusted to pH 6.0, 5.0, 4.5, & 4.0, and (F) selected *S. mitis* CMs or adjusted to pH7.0. One-way ANOVA was used to compare the mean of untreated control (#) or mch (*) with the mean of every other group and Dunnett post-test was used for multiple comparisons. Only significant relationships are shown (n=4, *p < 0.05, and ****p < 0.0001).

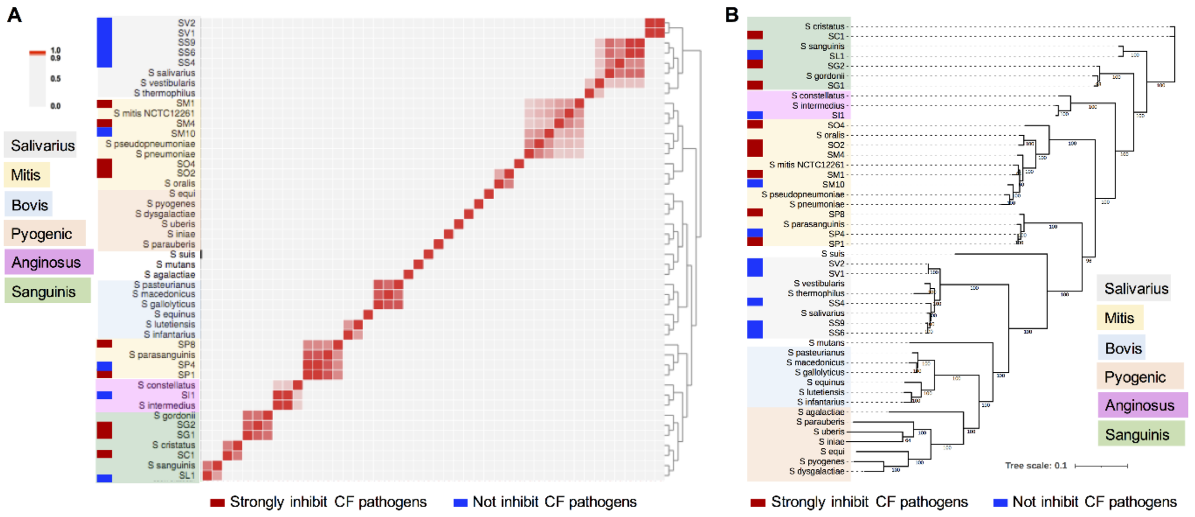


**Figure S4: Taxonomy investigation of *Streptococcus* CF isolates through ANI analysis and phylogenomic analysis using complete whole genome sequences.** A. Pairwise average nucleotide identity (ANI) analysis of complete whole genome sequences of the 18 CF isolates with 28 *Streptococcus* type strains. CF isolates showing inhibitory effects (SP8, SP1, SM1, SM4, SO2, SO4, SG2, SG1, SC1) are indicated by red bars, while CF isolates without inhibitory effects (SV2, SV1, SS4, SS9, SS6, SP4, SM10, SI1, SL1) are indicated by blue bars. Heatmap and dendrogram of ANI between these genomes are shown. The major *Streptococcus* groups Salivarius, Mitis, Bovis, Progenic, Anginosus, and Sanguinis are colour shaded. B. Phylogeny of the genomes of the 18 CF isolates and 28 *Streptococcus* type strains. The phylogenetic analysis was performed using the Maximum Likelihood method based on concatenated single-copy core genes and presented as an unrooted tree. The numbers indicate bootstrap values of the branches. The major *Streptococcus* groups are colour shaded.

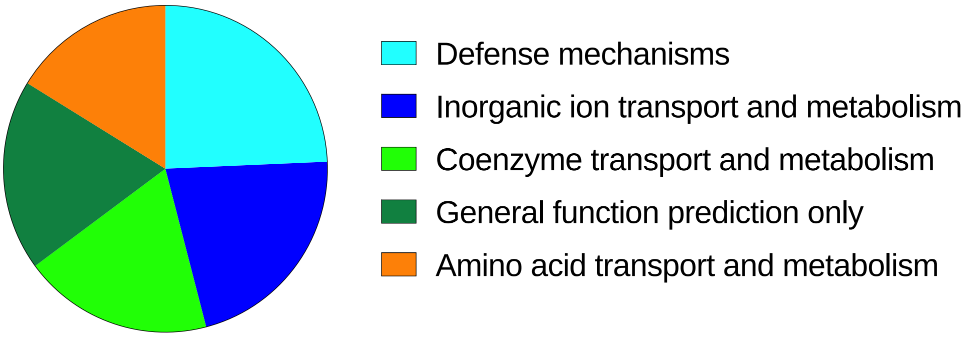


**Figure S5: Top five function categories highly enriched in the negative group that could not inhibit pathogen growth**. The relative ratios of the numbers of functions in each category are displayed as a pie chart.

#### 2.2 Supplementary Tables

**Table S1: Overview of the aerobic or facultative anaerobic commensal strains isolated from cystic fibrosis microbiome samples.**

| Genus | Species | No. of strains of each species |
|---|---|---|
| ***Streptococcus*** | *S. cristatus* | 3 |
| | *S. gordonii* | 3 |
| | *S. intermedius* | 1 |
| | *S. constellatus* | 1 |
| | *S. mitis* | 11 |
| | *S. sanguinis* | 2 |
| | *S. oralis* | 10 |
| | *S. parasanguinis* | 8 |
| | *S. salivarius* | 9 |
| | *S. vestibularis* | 3 |
| ***Neisseria*** | *N. flavescens* | 5 |
| | *N. macacae* | 4 |
| | *N. perflava* | 4 |
| | *N. subflava* | 7 |
| ***Actinomyces*** | *A. oris* | 4 |
| ***Corynebacterium*** | *C. durum* | 1 |
| | *C. pseudodiphtheriticum* | 1 |
| ***Dermabacter*** | *D. hominis* | 1 |
| ***Micrococcus*** | *M. luteus* | 2 |
| ***Rothia*** | *R. dentocariosa* | 2 |
| | *R. mucilaginosa* | 1 |

**Table S2. Overview of CF and non-CF pathogenic strains used in this study.**

| Genus | Species | Strain name | Reference |
|---|---|---|---|
| ***Achromobacter*** | | | |
| | *A. xylosoxidans* | AX1 | This study |
| ***Burkholderia*** | | | |
| | *B. multivorans* | BM1 | This study |
| ***Enterococcus*** | | | |
| | *E. faecalis* | ETF1 | This study |
| | | ATCC 27853 | ATCC 27853 |
| ***Haemophilus*** | | | |
| | *H. influenza* | ATCC 49247 | ATCC 49247 |
| ***Klebsiella*** | | | |
| | *K. pneumoniae* | KP1 | This study |
| | | ATCC 13883 | ATCC 13883 |
| ***Proteus*** | | | |
| | *P. mirabilis* | PTM1 | This study |
| ***Pseudomonas*** | | | |
| | *P. aeruginosa* | PAO1-mcherry | (Belon et al., 2015) |
| | | PADD1 | This study |
| | | PADD2 | This study |
| | | PADD3 | This study |
| | | PA8 | (Kolbe et al., 2020) |
| | | PA24 | (Kolbe et al., 2020) |
| | | CHA | (Dacheux, Attree, Schneider, & Toussaint, 1999; Toussaint, Delic-Attree, & Vignais, 1993) |
| | | ATCC 27853 | ATCC27853 |
| ***Staphylococcus*** | | | |
| | *S. aureus* | SPA4 | This study |
| | | ATCC 25923 | ATCC 25923 |
| ***Stenotrophomonas*** | | | |
| | *S. maltophilia* | STM1 | This study |

**Table S3: Genome sequencing statistics.**

| Isolate | coverage | # contigs | Largest contig | Total length | GC (%) | N50 | N75 | L50 | L75 |
|---|---|---|---|---|---|---|---|---|---|
| SC1 | 48 | 11 | 541575 | 1973970 | 42.45 | 496308 | 223341 | 2 | 4 |
| SG1 | 51 | 18 | 304562 | 2187339 | 40.31 | 185907 | 132417 | 5 | 8 |
| SG2 | 75 | 9 | 804400 | 2156808 | 40.46 | 653923 | 266487 | 2 | 3 |
| SI1 | 83 | 10 | 432635 | 1948549 | 37.56 | 331348 | 171079 | 3 | 5 |
| SN3(SL1) | 28 | 25 | 852688 | 2988825 | 42.08 | 475937 | 137723 | 3 | 6 |
| SM1 | 110 | 20 | 552068 | 2022463 | 39.76 | 214308 | 135863 | 3 | 6 |
| SPS1(SM10) | 197 | 30 | 361841 | 1981757 | 39.89 | 125014 | 90127 | 5 | 9 |
| SM4 | 136 | 33 | 245592 | 1874967 | 40.28 | 124961 | 83053 | 6 | 10 |
| SO2 | 26 | 45 | 528925 | 2509326 | 40.78 | 148540 | 59824 | 4 | 11 |
| SO4 | 25 | 35 | 263256 | 2062892 | 39.33 | 141077 | 81541 | 6 | 11 |
| SP1 | 20 | 41 | 228529 | 2183246 | 41.61 | 115765 | 59881 | 7 | 13 |
| SP4 | 29 | 49 | 204407 | 2185058 | 41.62 | 77327 | 41373 | 9 | 19 |
| SP8 | 29 | 28 | 670517 | 2224732 | 41.55 | 104393 | 59683 | 5 | 13 |
| SS4 | 19 | 20 | 426330 | 2392142 | 39.31 | 196889 | 162868 | 5 | 9 |
| SS6 | 22 | 45 | 193120 | 2271528 | 39.59 | 88895 | 52450 | 9 | 17 |
| SS9 | 21 | 41 | 182818 | 2230399 | 39.69 | 105651 | 57155 | 9 | 17 |
| SV1 | 28 | 36 | 270963 | 2092716 | 39.86 | 137892 | 54560 | 6 | 12 |
| SV2 | 24 | 34 | 258610 | 2090545 | 39.86 | 137892 | 51373 | 6 | 13 |

**Table S4. Mean pH values of some bacteria conditioned medium (CM)**

| Species | Conditioned medium | Mean pH |
|---|---|---|
| Ctrl | Columbia broth | 7.4 |
| *P. aeruginosa*-mcherry | mch CM | 7.69 |
| *S. mitis* | SM1 CM | 5.02 |
| | SM3 CM | 5.24 |
| | SM4 CM | 4.95 |
| | SM10 CM | 5.05 |
| *S. oralis* | SO2 CM | 4.78 |
| | SO3 CM | 4.73 |
| | SO5 CM | 4.91 |
| | SO9 CM | 5.04 |
| *S. cristatus* | SC1 CM | 5.44 |
| | SC2 CM | 5.51 |
| | SC3 CM | 5.16 |
| *S. constellatus* | SL1 CM | 4.83 |
| *S. intermedius* | SI1 CM | 4.72 |
| *N. perflava* | NP1 CM | 7.71 |
| *N. flavescens* | NF1 CM | 7.57 |
| *N. subtilis* | NS1 CM | 7.58 |
| *N. macacae* | NM1 CM | 7.41 |
| *A. oris* | ACO1 CM | 6.94 |
| | ACO3 CM | 4.67 |
| *B. licheniformis* | BL1 CM | 7.79 |
| *C. durum* | CBD1 CM | 7.94 |
| *C. pseudodiphtheriticum* | CBP1 CM | 7.69 |
| *D. hominis* | DH1 CM | 7.09 |
| *M. luteus* | ML1 CM | 7.82 |
| *R. dentocariosa* | RD1 CM | 5.28 |
| *R. mucilaginosa* | RM1 CM | 5.16 |

**Table S5. Genome sequence information of *Streptococcus* type strains.**

| Organism_name | Strain_name | Assembly_accession | Bioproject | Biosample |
|---|---|---|---|---|
| ***Streptococcus salivarius*** | strain=NCTC 8618 | GCF_000785515.1 | PRJNA224116 | SAMN03174835 |
| ***Streptococcus sanguinis*** | strain=NCTC7863 | GCF_900475505.1 | PRJNA224116 | SAMEA3672886 |
| ***Streptococcus thermophilus*** | strain=ATCC 19258 | GCF_010120595.1 | PRJNA224116 | SAMN11175069 |
| ***Streptococcus mutans*** | strain=NBRC 13955 | GCF_006739205.1 | PRJNA224116 | SAMD00169830 |
| ***Streptococcus pneumoniae*** | strain=NCTC7465 | GCF_001457635.1 | PRJNA224116 | SAMEA2479568 |
| ***Streptococcus pyogenes*** | strain=NCTC8198 | GCF_002055535.1 | PRJNA224116 | SAMEA2479569 |
| ***Streptococcus intermedius*** | strain=NCTC11324 | GCF_900475975.1 | PRJNA224116 | SAMEA4012327 |
| ***Streptococcus uberis*** | strain=NCTC3858 | GCF_900475595.1 | PRJNA224116 | SAMEA3871780 |
| ***Streptococcus lutetiensis*** | strain=NCTC13774 | GCF_900475675.1 | PRJNA224116 | SAMEA3905384 |
| ***Streptococcus gallolyticus*** | strain=NCTC13773 | GCF_900475715.1 | PRJNA224116 | SAMEA3881065 |
| ***Streptococcus oralis*** | strain=NCTC 11427 | GCF_900637025.1 | PRJNA224116 | SAMEA3936797 |
| ***Streptococcus parasanguinis*** | strain=ATCC 15912 | GCF_000164675.2 | PRJNA224116 | SAMN00113608 |
| ***Streptococcus parauberis*** | strain=NCFD 2020 | GCF_000187935.1 | PRJNA224116 | SAMN02436555 |
| ***Streptococcus equi*** | strain=ATCC 33398 | GCF_900156215.1 | PRJNA224116 | SAMN05421817 |
| ***Streptococcus infantarius subsp. infantarius*** | strain=ATCC BAA-102 | GCF_000154985.1 | PRJNA224116 | SAMN00000017 |
| ***Streptococcus equinus*** | strain=ATCC 9812 | GCF_000187265.1 | PRJNA224116 | SAMN00217012 |
| ***Streptococcus agalactiae*** | strain=ATCC 13813 | GCF_000186445.1 | PRJNA224116 | SAMN00217013 |
| ***Streptococcus pseudopneumoniae*** | strain=CCUG 49455 | GCF_002087075.1 | PRJNA224116 | SAMN06459226 |
| ***Streptococcus dysgalactiae*** | strain=NCTC6403 | GCF_901543725.1 | PRJNA224116 | SAMEA3649039 |
| ***Streptococcus pasteurianus*** | strain=NCTC13784 | GCF_900478025.1 | PRJNA224116 | SAMEA4030747 |
| ***Streptococcus gordonii*** | strain=Challis substr. CH1 | GCF_000017005.1 | PRJNA224116 | SAMN02603977 |
| ***Streptococcus macedonicus*** | strain=ACA-DC 198 | GCF_000283635.1 | PRJNA224116 | SAMEA2272145 |
| ***Streptococcus iniae*** | strain=YSFST01-82 | GCF_000831485.1 | PRJNA224116 | SAMN03286870 |
| ***Streptococcus suis*** | strain=S735 | GCF_000294495.1 | PRJNA224116 | SAMN02604110 |
| ***Streptococcus mitis*** | strain=NCTC12261 | GCF_000148585.2 | PRJNA173 | SAMN02435817 |
| ***Streptococcus vestibularis*** | strain=NCTC12167 | GCF_900636445.1 | PRJNA224116 | SAMEA3594358 |
| ***Streptococcus constellatus*** | strain=FDAARGOS_1156 | GCF_016725005.1 | PRJNA224116 | SAMN16357325 |
| ***Streptococcus cristatus*** | strain=ATCC51100 | GCF_900475445.1 | PRJNA224116 | SAMEA3649037 |

**Table S6: Functions highly enriched in the positive group.**

| FUNCTION (as protein name) | p_value | COG_ ACC | p_ yes | p_ no | COG_ PATHWAY | COG_ CATEGORY |
|---|---|---|---|---|---|---|
| Alpha-L-fucosidase (AfuC) (PDB:1HL8) | 1,48E+12 | COG3669 | 1,0 | 0,1 | | **Carbohydrate transport and metabolism** |
| Putative alpha-1,2-mannosidase (PDB:2WVX) | 7,13E+10 | COG3537 | 1,0 | 0,2 | | |
| Alpha-mannosidase (MngB) (PDB:5KBP) | 7,13E+10 | COG0383 | 1,0 | 0,2 | | |
| ABC-type polysaccharide transport system, permease component (LplB) (PDB:4TQU) | 0,00270 | COG4209 | 1,0 | 0,3 | | |
| N-acetyl-beta-hexosaminidase (Chb) (PDB:3RPM) | 0,00270 | COG3525 | 1,0 | 0,3 | | |
| Endo-beta-N-acetylglucosaminidase D | 0,00442 | COG4724 | 0,9 | 0,2 | | |
| Putative N-acetylmannosamine-6-phosphate epimerase (NanE) (PDB:1Y0E) | 0,00851 | COG3010 | 1,0 | 0,4 | | |
| 6-phosphogluconate dehydrogenase (Gnd) (PDB:2ZYA) | 0,00851 | COG0362 | 1,0 | 0,4 | Pentose phosphate pathway | |
| Phosphotransferase system, mannose/fructose/ N-acetylgalactosamine-specific component IIB (AgaB) (PDB:1BLE) | 0,00851 | COG3444 | 1,0 | 0,4 | | |
| Phosphotransferase system, mannose/fructose-specific component IIA (ManX) (PDB:1PDO) | 0,00851 | COG2893 | 1,0 | 0,4 | | |
| ABC-type glycerol-3-phosphate transport system, permease component (UgpE) (PDB:1VR4) | 0,00851 | COG0395 | 1,0 | 0,4 | | |
| Beta-galactosidase GanA (GanA) (PDB:1KWG) | 0,00851 | COG1874 | 1,0 | 0,4 | | |
| Ribose 5-phosphate isomerase RpiB (RpiB) (PDB:1NN4) | 0,00851 | COG0698 | 1,0 | 0,4 | Pentose phosphate pathway | |
| 6-phosphogluconolactonase, cycloisomerase 2 family (Pgl) (PDB:1JOF) | 0,00851 | COG2706 | 1,0 | 0,4 | | |
| Phosphoenolpyruvate synthase/pyruvate phosphate dikinase (PpsA) (PDB:5HV6) | 0,00851 | COG0574 | 1,0 | 0,4 | Gluconeogenesis | |
| Maltodextrin utilization protein YvdJ (function unknown) (YvdJ) | 0,00851 | COG5521 | 1,0 | 0,4 | | |
| ABC-type glycerol-3-phosphate transport system, periplasmic component (UgpB) (PDB:2Z8D) | 0,00851 | COG1653 | 1,0 | 0,4 | | |
| Beta-galactosidase, beta subunit (EbgC) (PDB:1JOP) | 0,00851 | COG2731 | 1,0 | 0,4 | | |
| Phosphotransferase system, galactitol-specific IIC component (SgcC) | 0,00851 | COG3775 | 1,0 | 0,4 | | |
| Glucose-6-phosphate 1-dehydrogenase (Zwf)(PDB:5B7W) | 0,00851 | COG0364 | 1,0 | 0,4 | Pentose phosphate pathway | |
| Phosphotransferase system, galactitol-specific IIB component (SgaB) (PDB:5GQS) | 0,00851 | COG3414 | 1,0 | 0,4 | | |
| Kojibiose phosphorylase YcjT (ATH1) (PDB:1H54) | 0,02335 | COG1554 | 0,4 | 0,0 | | |

| | | | | | | |
|---|---|---|---|---|---|---|
| Meiotically up-regulated gene 157 (Mug157) protein (function unknown) (PDB:2NVP) | 7,13E+10 | COG3538 | 1,0 | 0,2 | | **Function unknown** |
| Uncharacterized conserved protein, contains DUF2130 domain | 0,00851 | COG4487 | 1,0 | 0,4 | | |
| Uncharacterized conserved protein YycO, NlpC/P60 family (YycO) | 0,00851 | COG3863 | 1,0 | 0,4 | | |
| Uncharacterized membrane protein YesL (YesL) | 0,00851 | COG5578 | 1,0 | 0,4 | | |
| Uncharacterized conserved protein YsxB, DUF464 family (YsxB) (PDB:1S12) | 0,00851 | COG2868 | 1,0 | 0,4 | | |
| Uncharacterized membrane protein YjjP, DUF1212 family (YjjP) | 0,00851 | COG2966 | 1,0 | 0,4 | | |
| Uncharacterized membrane protein YjjB, DUF3815 family (YjjB) | 0,00851 | COG3610 | 1,0 | 0,4 | | |
| Agmatine/peptidylarginine deiminase (AguA) (PDB:1VKP) | 0,01562 | COG2957 | 0,7 | 0,1 | | **Amino acid transport and metabolism** |
| Spermidine synthase (polyamine aminopropyltransferase) (SpeE) (PDB:6BQ2) | 0,02335 | COG0421 | 0,4 | 0,0 | | |
| Saccharopine dehydrogenase, NADP-dependent (Lys9) (PDB:1E5L) | 0,02335 | COG1748 | 0,4 | 0,0 | Lysine biosynthesis | |
| Arginine/lysine/ornithine decarboxylase (LdcC) (PDB:1C4K) | 0,02335 | COG1982 | 0,4 | 0,0 | | |
| Arginine utilization protein RocB (RocB) | 0,04548 | COG4187 | 0,6 | 0,1 | | |
| Muramoyltetrapeptide carboxypeptidase LdcA (peptidoglycan recycling) (LdcA) (PDB:3TLZ) | 0,00851 | COG1619 | 1,0 | 0,4 | | **Cellwall/membrane/envelope biogenesis** |
| Cell shape-determining protein MreD (MreD) | 0,00851 | COG2891 | 1,0 | 0,4 | | |
| Fructoselysine-6-P-deglycase FrlB or related protein, duplicated sugar isomerase (SIS) domain (AgaS) (PDB:3C3J) | 0,00851 | COG2222 | 1,0 | 0,4 | | |
| ABC-type nitrate/sulfonate/bicarbonate transport system, periplasmic component (TauA) (PDB:2X26) | 0,00270 | COG0715 | 1,0 | 0,3 | | **Inorganic ion transport and metabolism** |
| ABC-type nitrate/sulfonate/bicarbonate transport system, ATPase component (TauB) | 0,00270 | COG1116 | 1,0 | 0,3 | | |
| ABC-type nitrate/sulfonate/bicarbonate transport system, permease component (TauC) | 0,00270 | COG0600 | 1,0 | 0,3 | | |
| Zn-dependent glyoxalase, PhnB family (PhnB) (PDB:5KEF) | 0,00270 | COG2764 | 1,0 | 0,3 | | **Energy production and conversion** |
| Glycerol kinase (GlpK) (PDB:2ZF5) | 0,04548 | COG0554 | 0,9 | 0,4 | | |
| DNA-binding transcriptional regulator, FrmR family (FrmR) (PDB:2HH7) | 0,02335 | COG1937 | 0,4 | 0,0 | | **Transcription** |

| | | | | | | |
|---|---|---|---|---|---|---|
| DNA-binding transcriptional regulator, PucR/PutR family (PucR) | 0,04548 | COG2508 | 0,9 | 0,4 | | |
| Diadenosine tetraphosphatase ApaH/serine/ threonine protein phosphatase, PP2A family (ApaH) (PDB:1AUI) | 0,00851 | COG0639 | 1,0 | 0,4 | | **Signal transduction mechanisms\|Transcription** |
| Two-component response regulator, YesN/ AraC family, consists of REC and AraC-type DNA-binding domains (YesN) | 0,00851 | COG4753 | 1,0 | 0,4 | | |
| Membrane protease subunit, stomatin/prohibitin family, contains C-terminal Zn-ribbon domain (YdjI) | 0,00442 | COG4260 | 0,8 | 0,1 | | **Posttranslational modification, protein turnover, chaperones** |
| Peroxiredoxin (Bcp) (PDB:5IPH) | 0,00851 | COG1225 | 1,0 | 0,4 | | |
| Thiamin-binding stress-response protein YqgV, UPF0045 family (YqgV) (PDB:1LXJ) | 0,00442 | COG0011 | 0,8 | 0,1 | | **Coenzyme transport and metabolism** |
| Nicotinamide riboside transporter PnuC (PnuC) (PDB:4QTN) | 0,00851 | COG3201 | 1,0 | 0,4 | | |
| HD superfamily phosphohydrolase (YdhJ) (PDB:2HEK) | 0,00851 | COG1078 | 1,0 | 0,4 | | **General function prediction only** |
| Predicted lipid-binding transport protein, Tim44 family (Tim44) (PDB:2CW9) | 0,00851 | COG4395 | 0,6 | 0,0 | | **Lipid transport and metabolism** |
| IMP dehydrogenase/GMP reductase (GuaB) (PDB:1B3O) | 0,00270 | COG0516 | 1,0 | 0,3 | Purine biosynthesis | **Nucleotide transport and metabolism** |

**Table S7: Functions highly enriched in the negative group.**

| FUNCTION (as protein name) | p_value | COG_ ACC | p_ yes | p_ no | COG_ PATHWAY | COG_ CATEGORY |
|---|---|---|---|---|---|---|
| CRISPR-Cas system-associated integrase Cas1 (Cas1) (PDB:2YZS) | 0,00270 | COG1518 | 0,0 | 0,7 | CRISPR-Cas system | **Defense mechanisms** |
| Alkyl hydroperoxide reductase subunit AhpC (peroxiredoxin) (AhpC) (PDB:2RII) | 0,00270 | COG0450 | 0,0 | 0,7 | | |
| Alkyl hydroperoxide reductase subunit AhpF (AhpF) (PDB:4YKG) | 0,00270 | COG3634 | 0,0 | 0,7 | | |
| CRISPR/Cas system endoribonuclease Cas6, RAMP superfamily (Cas6a) | 0,00851 | COG5551 | 0,0 | 0,6 | CRISPR-Cas system | |
| Abortive phage infection protein AbiEi, antitoxin component of an AbiEi-AbiEii TA system (AbiEi) | 0,00851 | COG5340 | 0,0 | 0,6 | | |
| Predicted nucleotidyltransferase component of viral defense system (PDB:4O8S) | 0,00851 | COG2253 | 0,0 | 0,6 | | |
| CRISPR/Cas system-associated endoribonuclease Cas2 (Cas2) (PDB:2I0X) | 0,00851 | COG1343 | 0,0 | 0,6 | CRISPR-Cas system | |
| Bacterial immunity/signal transduction membrane protein SdpI, SdpI/YbgB/DUF1648 family (SdpI) | 0,02335 | COG5658 | 0,6 | 1,0 | | |
| Antitoxin component HicB of the HicAB toxin-antitoxin system (HicB) (PDB:4P78) | 0,04548 | COG1598 | 0,1 | 0,6 | | |
| ABC-type enterochelin transport system, periplasmic component (CeuA) (PDB:3ZKW) | 7,13E+11 | COG4607 | 0,0 | 0,8 | | **Inorganic ion transport and metabolism** |
| Fe2+ transporter FeoB (FeoB) (PDB:3TAH) | 0,00442 | COG0370 | 0,1 | 0,8 | | |
| Fe2+ transport protein FeoA (FeoA) (PDB:2GCX) | 0,00442 | COG1918 | 0,1 | 0,8 | | |
| Copper chaperone CopZ (CopZ) (PDB:1AFI) | 0,00851 | COG2608 | 0,0 | 0,6 | | |
| Ammonia channel protein AmtB (AmtB) (PDB:1U77) | 0,00851 | COG0004 | 0,0 | 0,6 | | |
| ABC-type Co2+ transport system, permease component (CbiM) (PDB:4M58) | 0,01841 | COG0310 | 0,2 | 0,8 | | |
| Sulfate permease or related transporter, MFS superfamily (SUL1) (PDB:3NY7) | 0,02335 | COG0659 | 0,0 | 0,4 | | |
| Carbonic anhydrase (Cah) (PDB:1KOP) | 0,00851 | COG3338 | 0,0 | 0,6 | | |
| Ketopantoate reductase (PanE) (PDB:1BG6) | 0,00442 | COG1893 | 0,2 | 0,9 | Pantothenate/CoA biosynthesis | **Coenzyme transport and metabolism** |
| 7,8-dihydro-6-hydroxymethylpterin pyrophosphokinase (folate biosynthesis) (FolK) (PDB:1CBK) | 0,00851 | COG0801 | 0,0 | 0,6 | Folate biosynthesis | |
| Thiamine transporter ThiT (ThiT) (PDB:4POP) | 0,00851 | COG3859 | 0,0 | 0,6 | | |
| Phosphoglycerate dehydrogenase or related dehydrogenase (SerA) (PDB:3OET) | 0,01562 | COG0111 | 0,1 | 0,7 | Serine biosynthesis | |
| Phosphopantetheinyl transferase (Sfp) (PDB:1QR0) | 0,02335 | COG2091 | 0,0 | 0,4 | | |
| vWFA (von Willebrand factor type A) domain of Mg and Co chelatases (ChlD) | 0,04548 | COG1240 | 0,4 | 0,9 | | |
| Dihydroneopterin aldolase (FolB) (PDB:1B9L) | 0,00851 | COG1539 | 0,0 | 0,6 | Folate biosynthesis | |

| | | | | | | |
|---|---|---|---|---|---|---|
| Predicted epimerase YddE/YHI9, PhzF superfamily (YHI9) (PDB:1SDJ) | 0,00851 | COG0384 | 0,0 | 0,6 | | **General function prediction only** |
| Uncharacterized OsmC-related protein (YhfA) (PDB:1LQL) | 0,00851 | COG1765 | 0,0 | 0,6 | | |
| Uncharacterized membrane protein YozV, TM2 domain, contains pTyr (TM2) | 0,00851 | COG2314 | 0,0 | 0,6 | | |
| Predicted Na+-dependent transporter YfeH (YfeH) (PDB:4N7X) | 0,00851 | COG0385 | 0,0 | 0,6 | | |
| Uncharacterized membrane protein YedE/YeeE, contains two sulfur transport domains (YedE) | 0,02335 | COG2391 | 0,0 | 0,4 | | |
| Fatty acid repression mutant protein (predicted oxidoreductase) (FMR2) (PDB:2WQF) | 0,04548 | COG3560 | 0,1 | 0,6 | | |
| Urease beta subunit (UreB) | 0,00851 | COG0832 | 0,0 | 0,6 | | **Amino acid transport and metabolism** |
| Urease alpha subunit (UreC) (PDB:5A6T) | 0,00851 | COG0804 | 0,0 | 0,6 | | |
| Urease gamma subunit (UreA) (PDB:1A5K) | 0,00851 | COG0831 | 0,0 | 0,6 | | |
| Amino acid permease (LysP) | 0,00851 | COG0833 | 0,0 | 0,6 | | |
| Homocysteine/selenocysteine methylase (S-methylmethionine-dependent) (MHT1) (PDB:5DML) | 0,01562 | COG2040 | 0,1 | 0,7 | | |
| Small neutral amino acid transporter SnatA, MarC family (MarC) | 0,04548 | COG2095 | 0,1 | 0,6 | | |
| Beta-galactosidase/beta-glucuronidase (LacZ) (PDB:5N6U) | 0,00851 | COG3250 | 0,0 | 0,6 | | **Carbohydrate transport and metabolism** |
| Endo-1,4-beta-D-glucanase Y (BcsZ) (PDB:1CEM) | 0,01841 | COG3405 | 0,2 | 0,8 | | |
| Dihydroxyacetone kinase (DAK1) (PDB:1UN8) | 0,01841 | COG2376 | 0,2 | 0,8 | Isoprenoid biosynthesis | |
| Phosphotransferase system cellobiose-specific component IIC (CelB) (PDB:3QNQ) | 0,02335 | COG1455 | 0,6 | 1,0 | | |
| Beta-phosphoglucomutase, HAD superfamily (YcjU) (PDB:3E58) | 0,04548 | COG0637 | 0,4 | 0,9 | | |
| Hydrogenase/urease maturation factor HypB, Ni2+-binding GTPase (HypB)(PDB:2HF8) | 0,00851 | COG0378 | 0,0 | 0,6 | | **Posttranslational modification, protein turnover, chaperones** |
| Urease accessory protein UreE (UreE) (PDB:1EAR) | 0,00851 | COG2371 | 0,0 | 0,6 | | |
| Urease accessory protein UreH (UreH) (PDB:3SF5) | 0,00851 | COG0829 | 0,0 | 0,6 | | |
| Urease accessory protein UreF (UreF) (PDB:3SF5) | 0,00851 | COG0830 | 0,0 | 0,6 | | |
| Serine protease, subtilase family | 0,04548 | COG1572 | 0,1 | 0,6 | | |
| Cell fate regulator YmcA, YheA/YmcA/DUF963 family (controls sporulation, competence, biofilm development) (YmcA) (PDB:6PRH) | 0,00851 | COG4550 | 0,0 | 0,6 | | **Signal transduction mechanisms** |
| Cell fate regulator YaaT, PSP1 superfamily (controls sporulation, competence, biofilm development) (YaaT) | 0,00851 | COG1774 | 0,0 | 0,6 | | |
| Phosphotransferase subunit DhaM of the dihydroxyacetone kinase DhaKLM complex, contains PTS-EIIA, HPr, and PEP-utilizing domains (DhaM) | 0,01841 | COG3412 | 0,2 | 0,8 | | |
| Uncharacterized membrane protein YkvI (YkvI) | 0,00851 | COG3949 | 0,0 | 0,6 | | **Function unknown** |

| | | | | | | |
|---|---|---|---|---|---|---|
| Uncharacterized conserved protein, YiaAB domain (YiaAB) | 0,04548 | COG4298 | 0,1 | 0,6 | | |
| Predicted phospholipase, patatin/cPLA2 family (YjjU) | 0,00851 | COG4667 | 0,0 | 0,6 | | **Lipid transport and metabolism** |
| Acyl-CoA thioesterase FadM (FadM) (PDB:2FUJ) | 0,04548 | COG0824 | 0,1 | 0,6 | Menaquinone biosynthesis | |
| Nucleotide monophosphate nucleosidase PpnN/YdgH, Lonely Guy (LOG) family (PpnN)(PDB:2PMB) | 0,00851 | COG1611 | 0,0 | 0,6 | | **Nucleotide transport and metabolism** |
| Transposase InsO and inactivated derivatives (Tra5) | 0,01841 | COG2801 | 0,2 | 0,8 | | **Mobilome: prophages, transposons** |
| EndoRNase involved in mRNA decay, NYN (Nedd4-BP1/Rae1/YacP nuclease) family, contains PIN domain (Rae1) (PDB:5MQ8) | 0,01841 | COG3688 | 0,2 | 0,8 | | **Translation, ribosomal structure and biogenesis** |
| DNA repair exonuclease SbcCD nuclease subunit (SbcD) (PDB:6S85) | 0,00851 | COG0420 | 0,0 | 0,6 | | **Replication, recombination and repair** |
